# Aphid effector Mp10 balances immune suppression and defence activation through EDS1-dependent modulation of plant DAMP responses

**DOI:** 10.1101/2025.02.26.640334

**Authors:** Matteo Gravino, Sam T. Mugford, Daniela Pontiggia, Joshua Joyce, Claire Drurey, David C. Prince, Felice Cervone, Giulia De Lorenzo, Saskia A. Hogenhout

**Author notes:** Author for correspondence: *Saskia A. Hogenhout, Email:.

## Abstract

- Damage-associated molecular pattern (DAMP)-triggered immunity (DTI) serves as a crucial first line of defence against aphid attack, yet the mechanisms by which aphids suppress this response remain unclear.
- By investigating the colonisation of *Arabidopsis thaliana* by the highly polyphagous peach-potato aphid (*Myzus persicae*), we identified cell wall-derived DAMPs, specifically oligogalacturonides (OGs), as key in inducing DTI against aphids. The OG-responsive immune components BAK1/BKK1, CPK5/CPK6, GRP3, and EDS1 collectively contribute to DTI limiting aphid colonisation.
- We found that aphids limit OG production during feeding. Additionally, the salivary chemosensory protein (CSP) effector Mp10/CSP4, which is known to be delivered into the cytoplasm of plant cells early in aphid attack, inhibits OG-induced DTI.
- While Mp10 suppresses OG-induced DTI, it also interacts with EDS1-mediated defences, enhancing aphid fecundity in the absence of EDS1 but restoring OG responsiveness under attack, revealing its broader role in immune modulation and effector-driven host adaptation.

## INTRODUCTION

Aphids (Hemiptera: Aphididae) are highly specialised sap-feeding insects and major vectors of economically significant plant viruses. Their ability to colonise a wide range of plants depends on their feeding strategy and molecular interactions with the host. Using their piercing-sucking stylets, aphids navigate between plant cells, causing minimal tissue damage to reach the phloem sieve elements for sustained feeding (Tjallingii & Esch, 1993). However, this process involves repeated penetration of various plant cells, which can cause mechanical damage and trigger plant-defence responses. Plants detect such damage through the release of damage-associated molecular patterns (DAMPs) that activate DAMP-triggered immunity (DTI). Yet, aphids appear to suppress these responses to facilitate colonisation. Although aphid feeding has been linked to DTI (Silva-Sanzana *et al*., 2022), the underlying plant genes remain largely unknown, and the molecular mechanisms aphids use to counteract this defence are poorly understood.

Aphids not only cause direct damage but also play a major role in virus transmission. Viruses can be acquired from various plant tissues and transmitted either by attaching to the inner linings of the stylets and foregut or by circulating throughout the aphid body and being released *via* saliva (Whitfield *et al*., 2015). Aphids are particularly effective in this role, transmitting nearly 40% of all known vector-borne plant virus species (Peters *et al*., 2024). Given that most plant viruses rely on vectors for transmission (Ng & Falk, 2006; Hogenhout *et al*., 2008; Ammar *et al*., 2009), understanding plant defences against aphids is essential for developing effective pest and virus management strategies.

Plants have evolved multilayered immune responses to counteract herbivores and pathogens. Pattern-recognition receptors (PRRs) on the cell surface detect pathogen- and damage-associated molecular patterns (PAMPs/DAMPs), triggering PTI/DTI as an initial defence layer (Hou *et al*., 2019). However, successful pathogens and herbivores evade these defences by delivering effectors that suppress PTI/DTI. Intracellular nucleotide-binding leucine-rich repeat (NLR) receptors recognise these effectors and trigger a stronger effector-triggered immunity (ETI), often culminating in localised cell death to restrict further spread (Jones & Dangl, 2006). PTI and ETI function in an interconnected manner to provide full resistance (Yuan *et al*., 2021), with ENHANCED DISEASE SUSCEPTIBILITY 1 (EDS1) playing a key role in coordinating these immune responses (Dongus & Parker, 2021).

Damage perception is a critical component of plant immunity against herbivores (Gatehouse, 2002). Plant cell walls, primarily composed of polysaccharides such as cellulose, pectin, and hemicelluloses, are continuously monitored for structural integrity (Hamann & Denness, 2011; Molina *et al*., 2024). When cell wall-derived fragments are released, they act as DAMPs to activate DTI (Aziz *et al*., 2007; Claverie *et al*., 2018; De Lorenzo & Cervone, 2022). Consistent with this, aphid saliva contains enzymes, such as pectin methyl esterase (PME), polygalacturonase (PG), and cellulase (Cherqui, 2000; Ni *et al*., 2000; Guo *et al*., 2006), which can remodel plant cell walls and potentially promote the release of DAMPs (Molina *et al*., 2024; Pontiggia *et al*., 2024). Moreover, aphid feeding enhances plant PME and pectate lyase (PL) activity, leading to modifications in pectin structure (Silva-Sanzana *et al*., 2019). However, it remains unclear whether aphid-induced DAMPs are released in sufficient quantities to trigger DTI or how aphids manipulate this process to evade plant defences.

Aphids secrete two types of saliva: ‘sheath’ saliva, which forms a protective barrier around stylets, and ‘watery’ saliva, which is injected into cells during probing (Tjallingii & Esch, 1993). The latter type of saliva contains effectors that modulate plant immunity, including the salivary chemosensory protein (CSP) effector Mp10/CSP4, which is delivered into mesophyll cells during early feeding (Mugford *et al*., 2016). Notably, Mp10 suppresses PTI induced by bacterial flg22 and aphid-derived elicitors by interfering with plant deubiquitinating enzymes such as ASSOCIATED MOLECULE WITH THE SH3 DOMAIN OF STAM (AMSH), leading to destabilisation and mislocalization of PRRs (Bos *et al*., 2010; Drurey *et al*., 2019; Gravino *et al*., 2024). However, Mp10 also induces chlorosis with cell death-like features and reduces aphid reproduction (Bos *et al*., 2010; Rodriguez *et al*., 2014; Zhang *et al*., 2023; Rao *et al*., 2024). Mp10 is detected at aphid stylet tips (acrostyles) (Deshoux *et al*., 2022), enabling early delivery of effectors and viruses during probing and feeding (Uzest *et al*., 2007; Deshoux *et al*., 2022).

Importantly, aphid performance is negatively affected by treatment with oligogalacturonides (OGs), which are cell wall-derived DAMPs (Gravino, 2018; Silva-Sanzana *et al*., 2022). OGs activate immune-signalling pathways involving BRASSINOSTEROID INSENSITIVE 1-ASSOCIATED RECEPTOR KINASE 1 (BAK1) and its closest homolog BAK1-LIKE 1 (BKK1) co-receptors and calcium-dependent protein kinases (CDPKs), including CPK5 and CPK6 (Moscatiello *et al*., 2006; Gravino *et al*., 2015; Gravino *et al*., 2017), both of which have been implicated in plant-aphid interactions (Chaudhary *et al*., 2014; Prince *et al*., 2014a; Vincent *et al*., 2017). Moreover, GLYCINE-RICH PROTEIN 3 (GRP3) also plays a role in OG-mediated immune signalling (Gramegna *et al*., 2016). Notably, long OGs with a degree of polymerisation (DP) between 10 and 15 are the most active in triggering DTI, whereas short OGs (DP2-3) exhibit weaker immune activity and can even suppress PTI (Mathieu *et al*., 1991; Vorhölter *et al*., 2012; Gramegna *et al*., 2016; Davidsson *et al*., 2017; Xiao *et al*., 2024). Plants regulate OG signalling through specific OG oxidases that catalyse OG inactivation (Benedetti *et al*., 2018; Pontiggia *et al*., 2020; Salvati *et al*., 2024).

By investigating the colonisation of *Arabidopsis thaliana* by the highly polyphagous peach-potato aphid (*Myzus persicae*), we show that OG-induced DTI relies on BAK1/BKK1, CPK5/CPK6, GRP3, and EDS1 to restrict aphid colonisation. Aphids counteract this by limiting OG production and deploying Mp10, which inhibits OG and flg22 sensitivity and PRR stability while simultaneously inducing ETI-related defences. In EDS1-deficient plants, Mp10 enhances aphid fecundity, promotes PRR stability, and restores OG and flg22 responsiveness, highlighting a dual role in immune modulation. These findings reveal aphid strategies for evading plant immunity and align with recent discoveries that other aphid effectors actively target EDS1 (Liu *et al*., 2024), further illustrating the complexity of plant-aphid interactions.

## MATERIALS AND METHODS

### Plant materials

*A. thaliana* (hereafter Arabidopsis) ecotype Columbia-0 (Col-0) wild-type (WT) seeds were purchased from Lehle Seeds. *Nicotiana benthamiana* WT seeds were provided by JIC Horticultural Service. Transgenic lines used in this study include: Arabidopsis Col-0 *grp3* (SALK_084685.46.60) and *GRP3*-overexpressing (OE) lines (*35S:GRP3::RFP* #16-4) (Gramegna *et al*., 2016); Arabidopsis Col-0 *eds1-2* (Bartsch et al., 2006); Arabidopsis Col-0 *bak1-5 bkk1-1* (Schwessinger et al., 2011); Arabidopsis Col-0 *cpk5 cpk6* (Gravino *et al*., 2015), and *cpk5 cpk6 cpk11* (Boudsocq *et al*., 2010); Arabidopsis Col-0 OG-machine (OGM) β-oestradiol-inducible lines (*XVE:OGM*), and OGM lines under the *PATHOGENESIS-RELATED GENE 1* (*PR1*) promoter (*pPR1:OGM* #2) (Benedetti *et al*., 2015); *N. benthamiana eds1* (Schultink *et al*., 2017); *N. benthamiana adr1 nrg1* (Prautsch *et al*., 2023); *N. benthamiana nrg1-1* and *nrg1-2* (Qi *et al*., 2018); and *N. benthamiana* NahG (Wulff *et al*., 2004).

### Generation of Mp10 stable transgenic lines

For the generation of stable transgenic Arabidopsis lines expressing *Mp10* (amino acids 23–145) with an N-terminal Flag-tag in place of its signal peptide under a dexamethasone (DEX)-inducible promoter, *Mp10* was amplified from aphid cDNA using a forward primer that encodes a DYKDDDDK (Flag)-tag and a reverse primer carrying a stop codon (Supporting Information Table S1) and cloned into pBAV150 (Vinatzer *et al*., 2006) using Gateway technology (Invitrogen). Recombined plasmid was introduced into *Agrobacterium tumefaciens* strain GV3101-pMP90RK for subsequent transformation of Arabidopsis Col-0 WT using the floral dip method (Bechtold *et al*., 1993). Transgenic seeds were selected on phosphinothricin (BASTA). T2 seedlings exhibiting live versus dead segregation ratios of 3:1 were taken forward to the T3 stage, and two T3 lines (#7-5 and #9-5) with a 100% survival ratio (considered homozygous) were selected for experiments.

The *eds1-2* x Mp10 #7-5 and *eds1-2* x Mp10 #9-5 plants were generated *via* crossing the above-mentioned homozygous Mp10 #7-5 and #9-5 lines with the *eds1-2* mutant in the Col-0 background (Bartsch *et al*., 2006). Transgenic seeds were selected on BASTA. T3 seedlings with a 100% survival ratio were selected and genotyped by PCR (Chang *et al*., 2019) (Supporting Information Table S1) to isolate the T3 lines harbouring the *Mp10* and the *eds1-2* homozygous alleles.

For experiments with adult plants, Arabidopsis 1-week (w)-old seedlings were transplanted in 8 cm plastic pots (one seedling per pot) containing 0.4 l compost and grown in a controlled environment room (CER) under a 10 h, 22 °C : 14 h, 22 °C light/dark temperature regime and 70% humidity. Plants were used for assays 2 or 3 weeks (w) after transplanting.

### Aphids

Stock colonies of *M. persicae* clone O (Mathers *et al*., 2017) were maintained on Arabidopsis plants, ecotype Col-0 wild type (WT), in a CER under a 14 h, 24 °C : 10 h, 15°C light/dark temperature regime and 48% humidity. Age-synchronised aphids were used for the fecundity assays. To achieve this, adult female aphids were transferred from the stock colonies to Col-0 WT plants. After 24 h, the adult females were removed from the plants, while the progeny (age-synchronised nymphs) produced by the females were used for various assays.

### Isolation and analysis of OGs from aphid-infested Arabidopsis leaves

Arabidopsis Col-0 WT 4-w-old plants were infested with 6-d-old aphids (twenty or two aphids for isolation of OGs after 6 h or after 7/9 d of infestation, respectively) on the first fully expanded leaf from the top and caged inside a clip cage made with transparent plastic tubes and nylon mesh (Prince *et al*., 2014a; Prince *et al*., 2014b) to minimise the impact of the clip cage on leaf photosynthesis (Kou *et al*., 2022). After 6 h post infestation (hpi) or 7/9 d post infestation (dpi), leaves were excised from plants and cleaned from aphids using a paintbrush (ESPO). Two leaves per treatment were floated in ultrapure water in 6-well plates (Thermo Fisher Scientific) to remove any trace of insect debris. Controls comprised leaves exposed to empty clip cages and leaves without clip cages (untreated).

OGs were isolated with the leaf-strip method, as previously described (Benedetti *et al*., 2017). Briefly, leaf surfaces were sterilised for 3 min with 4 ml of 1% NaClO and washed four times with 6 ml of sterile ultrapure water. After removing the tip and the petiole, leaves were sliced in 2-mm-wide strips using a sterile scalpel blade (Slaughter Ltd, R & L) and incubated for 16 h at 30 °C with gentle shaking in 2 ml of strong chelating solution (i.e., 50 mM ammonium acetate pH 5.0, 50 mM CDTA, 50 mM ammonium oxalate, Sigma-Aldrich) supplemented with 10 mM sodium sulphite (Sigma-Aldrich) to preserve the OGs and inhibit the activity of OG oxidases during the extraction, or without sodium sulphite, for the eventual detection of oxidised OGs (Benedetti *et al*., 2018). The incubation medium was collected, diluted with 8 ml of 100% ethanol (VWR Chemicals), and centrifuged at 15,000 ***g*** for 30 min into an Oak Ridge polypropylene centrifuge tube (Thermo Fisher Scientific). After discarding the supernatant, the pellet was air-dried inside a laminar flow cabinet and resuspended in 100 μl of ultrapure water.

Peaks, including OG oligomers from DP6 to DP15, were separated and analysed by high-performance anion-exchange chromatography (HPAEC) with pulsed amperometric detection (PAD). Data were normalised using the compositional data normalisation (CoDA) method, as previously described (Aitchison, 1989; Noonan *et al*., 2018). Briefly, the abundance of OGs was calculated as the log-ratio of their peak area to the geometric mean of all peaks in the profile that were shared between different treatments, using the equation:

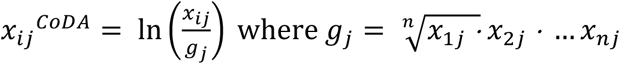

where *x_ij_^CoDA^* is the normalised peak area of the *i*th peak in the *j*th profile, *xij* is the *i*th peak area in the *j*th profile, and *g_j_* is the geometric mean of all peaks in the *j*th profile that were shared between different treatments. Because log-transformation of values lower than 1 generates negative values, transformed data were corrected by adding a constant value, a = 4, to obtain all positive values.

### Preparation of OGs for plant immunity assays

OGs enriched in DP10-15 were prepared from solutions of high molecular weight unmethylated polygalacturonic acid (Alfa Aesar) digested with an endo-PGII from *Aspergillus niger*, as previously described (Benedetti *et al*., 2017). OG powder was freshly dissolved in ultrapure water and diluted to working solutions.

### Quantitative reverse transcriptase (qRT)-PCR assays

Arabidopsis Col-0 WT and Mp10-inducible 3-to 4-w-old plants were sprayed with 1.25 μM DEX. After 3 d, plants were infiltrated with ultrapure water or OGs (200 μg ml^-1^) on the abaxial surface of the first fully expanded leaf from the top. After 2 h, infiltrated leaves were collected, and total RNA was extracted using Tri-Reagent (Sigma-Aldrich). Complementary DNA (cDNA) was synthesised from 250 ng of DNase I (RQ1 RNase-free DNase, Promega)-treated RNA using the reverse transcriptase M-MLV Kit (Invitrogen). Reactions of qRT-PCR, consisting of 2.5 ng cDNA, 1 μM of each primer (Supporting Information Table S1), 1X SYBR Green JumpStart Taq ReadyMix (Sigma-Aldrich), and nuclease-free water up to 10 μl, were performed in white 96-well plates (4titude) and detected by a CFX96 Real-Time System with a C1000 Thermal Cycler (Bio-Rad).

The expression levels of defence-related genes of interest (GOI), *FLG22-INDUCED RECEPTOR-LIKE KINASE 1* (*FRK1*, *At2g19190*), *CYTOCHROME P450, FAMILY 81, SUBFAMILY F, POLYPEPTIDE 2* (*CYP81F2*, *At5g57220*), *PHYTOALEXIN DEFICIENT 3* (*PAD3*, *At3g26830*) and *4* (*PAD4*, *At3g52430*) were normalised to those of housekeeping genes (HKGs) *GLYCERALDEHYDE-3-PHOSPHATE DEHYDROGENASE C2* (*GAPDH*, *At1g13440*) and *UBIQUITIN 5* (*UBQ5*, *At3g62250*) as previously described (Pfaffl, 2001; Vandesompele *et al*., 2002; Ferrari *et al*., 2006), with the following modifications. For each gene, the average threshold cycle (aCt) from three technical replicates per biological sample was corrected for the PCR efficiency (E) to obtain Ct using the equation aCT × log(E,2). The geometric mean (GM) of Ct values of HKGs was then calculated. The ΔCt between GOIs and HKGs was calculated as Ct_GOI_ -GM of Ct_HKGs_. The expression levels for each GOI were calculated as 2^−ΔCT^ to obtain gene expression values relative to *GAPDH* and *UBQ5*.

### Analysing aphid performance on OG-exposed plants

Arabidopsis WT and mutant/transgenic 4-w-old plants were infiltrated with OGs (200 μg ml^-1^) or ultrapure water, as a control, on the abaxial surface of the first fully expanded leaf from the top. After 72 h, 6-d-old age-synchronised adult aphids were placed on the infiltrated leaf using a moist paintbrush at one aphid in a clip cage per leaf (Prince *et al*., 2014a; Prince *et al*., 2014b). After 10 d, the numbers of aphids inside each clip cage were counted. For assays with Mp10-inducible lines, plants were sprayed with 1.25 μM DEX 3 d before elicitor treatment. Each experiment included ten plants per sample, unless otherwise stated.

### Aphid whole-plant fecundity assays

Arabidopsis Col-0 WT and transgenic 3-w-old plants were seeded with one 1-d-old age-synchronised nymph, individually caged in transparent plastic tubes (10-cm diameter, 30-cm height) capped with white gauze-covered plastic lids both on top and on the bottom. After *ca.* 6 d, the nymphs developed into adults capable of producing their own offspring, which were counted (and removed at each count) on days 7, 9, and 11 and added up to determine the total number of nymphs produced per adult. Each experiment included eight plants per genotype.

### Chlorophyll extraction and quantification in Arabidopsis

Chlorophyll extraction and quantification were performed as previously described (Sieber *et al*., 2000). Three-week-old Arabidopsis plants, including Col-0 WT, the *eds1-2* mutant, and Mp10-inducible in both Col-0 and *eds1-2* backgrounds, were sprayed twice with 1.25 μM DEX at 6-d intervals. After 12 d, 3-5 leaves (approximately 150 mg) were collected from each plant and placed in 15 ml tubes containing 10 ml of 80% ethanol. Tubes were covered with foil to protect them from light and rotated on a shaker.

Chlorophyll content was measured after 24 h of incubation using a NanoDrop 1000 spectrophotometer (Thermo Scientific) at 647 and 664 nm. The micromolar (µmol) concentration of total chlorophyll was calculated using the following equation and normalised to g of sample fresh weight:

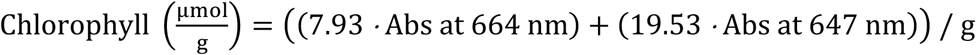

For dark-induced senescence, the plants were sprayed as described above and then incubated in complete darkness for 5 d before chlorophyll measurements. Whole plants (approximately 90 mg) were used, incubated in 5 ml tubes containing 4 ml of 80% ethanol, and processed similarly.

### Detection of hydrogen peroxide by DAB staining in Arabidopsis

Arabidopsis plants were grown and treated as described in the above section. Leaves were collected and placed in 6-well plates (Thermo Fisher Scientific) containing at least 4 ml of 3,3’-diaminobenzidine (DAB) staining solution (Daudi & O’Brien, 2012). *In situ* detection of hydrogen peroxide was performed as previously described (Daudi & O’Brien, 2012).

### Agrobacterium-mediated transient expression in *N. benthamiana*

*N. benthamiana* plants were grown in a CER under a 16 h, 22 °C : 8 h, 22 °C light/dark temperature regime and 80% humidity. The constructs containing eGFP-Mp10 or eGFP alone under the *Cauliflower mosaic virus* (CaMV) 35S promoter in pB7WGF2 or pB7WG2 plasmids (Karimi *et al*., 2002), respectively, were previously described (Gravino *et al*., 2024). The constructs containing Flag-Mp10 or Flag-alone under the CaMV 35S promoter in pJL-TRBO plasmid (Lindbo, 2007) were previously described (Gravino *et al*., 2024). The construct containing AtFLS2-3xMyc-eGFP under the *FLS2* promoter in pCAMBIA 2300 was previously described (Robatzek *et al*., 2006). The constructs were introduced into *A. tumefaciens* strain GV3101-pMP90RK, which was used for agroinfiltration on each side of the abaxial base of two expanded leaves of 4-to 5-w-old *N. benthamiana* plants at a final concentration of OD_600_=0.3-0.5 in MMA buffer (10 mM MES pH= 5.5, 10 mM MgCl2, 100 µM acetosyringone) as previously described (Bos *et al*., 2010). The Agrobacterium carrying the pCB301-p19 plasmid (Win & Kamoun, 2004) was supplemented at OD_600_=0.1. Leaf/plant assays were conducted 3-14 days post infiltration.

### Chlorosis assays in *N. benthamiana*

Chlorosis symptoms in the areas agroinfiltrated with eGFP-Mp10, but not eGFP alone, became visible at 3 days post-infiltration (dpi), and images were taken at 7 dpi. Chlorosis was quantified using Fiji (Schindelin *et al*., 2012) by selecting a region of interest (ROI) including the agroinfiltrated area. The ROI was analysed using a colour histogram to extract the red, green, and blue (RGB) components. The yellow component, indicative of chlorosis, was calculated using the formula: (0.5 × red) + (0.5 × green).

For systemic chlorosis and dwarfism phenotyping, Agrobacteria carrying either Flag-Mp10 or Flag alone were infiltrated into one side of a single leaf in different plants. Symptoms became visible around 10 dpi, and images were taken at 14 dpi.

### Protein extraction, immunoblotting, and antibodies

Three days after agroinfiltration, two leaf discs of 7 mm diameter were obtained from the agroinfiltrated area in at least three plants per construct mix (i.e., eGFP-AtFLS2 with Flag alone or Flag-Mp10) and ground into a fine powder using a TissueLyser LT Bead Mill (Qiagen). The powder was homogenised in 150 µl of protein extraction/loading buffer (10 mM Tris-HCl at pH 7.5, 50 mM dithiothreitol, 4% sodium dodecyl sulphate, 10% glycerol, 0.05% bromophenol blue), vortexed, boiled for 5 min, and centrifuged at 5,000 ***g*** for 3 min. Western blotting was performed as previously described (Gravino *et al*., 2024) using 10 µl of protein extract. Antibodies were as described (Gravino *et al*., 2024), except for anti-FLS2-rabbit (PhytoAB), which was diluted 1:2000.

### ROS burst assays

ROS production was determined using a luminol/peroxidase-based method as previously described (Savatin *et al*., 2014a; Gigli-Bisceglia *et al*., 2015), with some modifications. For ROS burst in Arabidopsis, Col-0 WT and Mp10-inducible 3-to 4-w-old plants were sprayed with 1.25 μM DEX. After 2 d, at least 24 leaf discs of 4 mm diameter were obtained from at least three plants for each genotype, incubated in white 96-well plates (Grenier Bio-One LUMITRAC^TM^) containing 200 μl of ultrapure water per well supplemented with 1.25 μM DEX, and covered with aluminium foil. After 1 d, the DEX solution was replaced with 100 μl of a solution containing 10 μg ml^-1^ horseradish peroxidase (Sigma-Aldrich) and 17 μg ml^-1^ luminol (Sigma-Aldrich) supplemented with flg22 peptide (QRLSTGSRINSAKDDAAGLQIA, 100 nM, EZBiolab), chitin oligosaccharide (500 μg ml^-1^, Yaizu Suisankagaku Industry, YSK, Japan), OGs (200 μg ml^-1^), or water, as control. OGs and controls were vacuum infiltrated for 2 m prior to imaging. Luminescence was captured and processed using a Photek camera (East Sussex, UK) and the IFS32 imaging software (Photek). Luminescence data were acquired every 30 s for at least 30 m.

For ROS burst in *N. benthamiana*, the same number and size of leaf discs was obtained from the agroinfiltrated area in at least two plants per construct (i.e., eGFP alone or eGFP-Mp10) and processed as described above.

### Statistical analyses

All statistical analyses have been performed in R packages (R Core Team, 2019) or JASP (JASP Team, 2024) using a Student’s *t*-test, one-way or two-way ANOVA with post-hoc Tukey HSD test. Experiments consisted of at least three biological replicates and were repeated at least two times on different days to generate data from at least two independent experiments, which were displayed as plots using R packages (R Core Team, 2019).

## RESULTS

### OG-triggered immunity reduces *M. persicae* fecundity on Arabidopsis *via* BAK1/BKK1-, CPK5/6-, and GRP3-mediated defence pathways

We examined whether plant immunity triggered by DAMPs, such as OGs, plays a role in plant defence against aphids. We found that the fecundity of clonally (asexually) reproducing *M. persicae* aphids was reduced on Arabidopsis Col-0 WT leaves infiltrated with elicitor-active exogenous OGs enriched in degree of polymerisation (DP) 10-15 (Benedetti *et al*., 2017) (Fig. 1a,b,c) compared to leaves infiltrated with water (Fig. 1b,c). We also measured aphid fecundity on transgenic Arabidopsis plants expressing a polygalacturonase-inhibiting protein (PGIP)-PG chimaera, referred to as the “OG machine” (OGM) (Fig. 1d; Supporting Information Fig. S1a). Upon treatment with 5 μM β-oestradiol dissolved in DMSO, these plants accumulated elicitor-active OGs *in vivo* (Benedetti *et al*., 2015), leading to the activation of DTI (Supporting Information Fig. S1a) and the reduction of aphid fecundity compared to plants treated with DMSO alone (Fig. 1b,e). The 5 μM β-oestradiol treatment itself did not affect aphid fecundity (Supporting Information Fig. S1b). It is well known that the expression of *PR1* is induced in Arabidopsis ecotypes such as Col-0 infested with *M. persicae* (Moran & Thompson, 2001; Kusnierczyk *et al*., 2007; Kettles *et al*., 2013). Hence, we used the *PR1* promoter to drive the OG production by the chimeric *OGM* gene (Benedetti *et al*., 2015). We found that aphid fecundity was reduced on *pPR1:OGM* plants (Fig. 1d), compared to Col-0 WT plants (Fig. 1f,g). These results indicate that *M. persicae* fecundity is negatively affected by OG-induced DTI, in agreement with previous findings (Gravino, 2018; Silva-Sanzana *et al*., 2022).

**Fig. 1.**
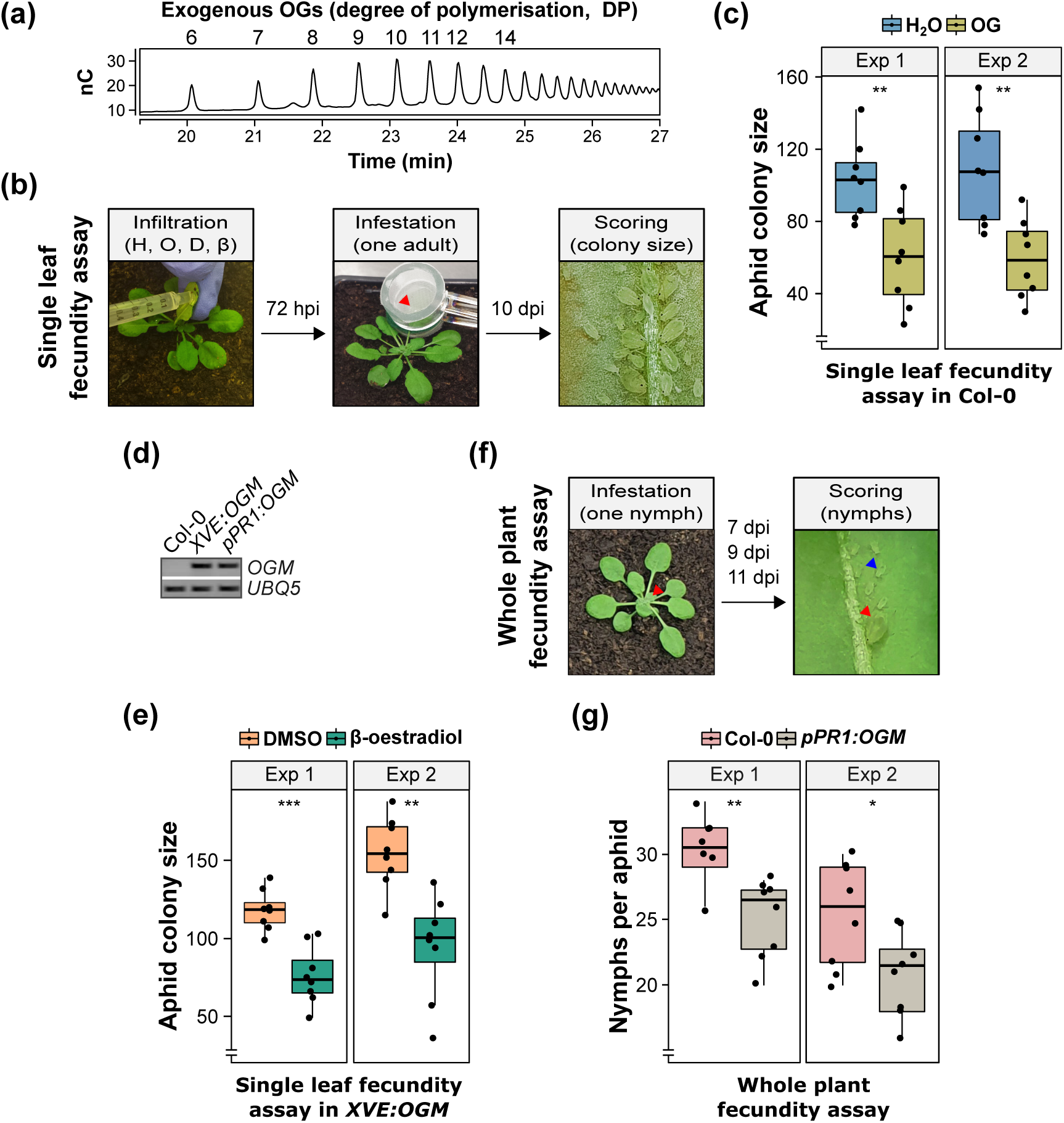
Exogenous and endogenous oligogalacturonides (OGs) increase Arabidopsis resistance against *Myzus persicae*. (a) Profile of exogenous OGs (2.5 μg) enriched in degree of polymerisation (DP) 10-15 used in this work, analysed by high-performance anion-exchange chromatography (HPAEC) with pulsed amperometric detector (PAD). The x-axis shows the retention time in min. The y-axis shows the detector response in nanocoulombs (nC). (b) Experimental design of aphid single-leaf fecundity assay. One leaf from 4-week (w)-old Arabidopsis Col-0 wild type (WT) or *XVE:OGM* transgenic plants was infiltrated with H_2_O (H) and 200 μg ml^-1^ OGs (O), or DMSO (D) and 5 μM β-oestradiol (β), respectively (left panel). 72 hours post infiltration (hpi), one 6-d-old asexually reproducing *M. persicae* adult female was caged on the infiltrated leaf (red arrow, middle panel). After 10 d, the number of individual aphids (adults + nymphs) within the cage was counted to obtain the aphid colony size (right panel). (c) Aphid single-leaf fecundity assay in Col-0 WT plants pre-treated with OGs or H_2_O as a control. The y-axis shows the aphid colony size. (d) Genotyping of *XVE:OGM* and *pPR1:OGM* transgenic plants using gene-specific primers listed in Supporting Information Table S1 and genomic DNA as a template. (e) Aphid single-leaf fecundity assay in *XVE:OGM* transgenic plants pre-treated with β- oestradiol or DMSO as a control. The y-axis shows the aphid colony size. (f) Experimental design of aphid whole plant fecundity assay. Arabidopsis Col-0 WT and *pPR1:OGM* transgenic 3-w-old plants were infested with one 1-d-old *M. persicae* nymph (red arrow, left panel). After 7, 9, and 11 dpi, the progeny (blue arrow, right panel) of this aphid (red arrow, right panel) was counted and removed from the plant and added up to obtain the number of nymphs produced per individual aphid. (g) Aphid whole plant fecundity assay in Col-0 WT and *pPR1:OGM* transgenic plants. The y-axis shows the number of nymphs per aphid. In (c, e, g), boxplots show the median, the 25^th^ and 75^th^ percentiles, the most extreme data points (whiskers’ extensions), and the observations as black filled circles. Two independent experiments (exp) are shown. n = 8 in each experiment. Asterisks indicate significant differences between samples as determined by Student’s *t* test (*, *P* < 0.05; **, *P* < 0.01; ***, *P* < 0.001).

DTI induction by OGs is dependent on key immune-related elements, including BAK1 and BKK1, CPK5 and CPK6, and GRP3 (Gravino *et al*., 2015; Gramegna *et al*., 2016; Gravino *et al*., 2017) (Supporting Information Table S2). Here we assessed if these immune-signalling components affect the OG-induced reduction of aphid fecundity on Arabidopsis. Because BAK1 and BKK1 play a redundant and equal contribution in OG immune signalling (Gravino *et al*., 2017), we assessed aphid performance on the *bak1-5 bkk1-1* double mutant (Schwessinger *et al*., 2011) (Fig. 2a; Supporting Information Fig. S2). Unlike Col-0 WT plants, aphid fecundity was not reduced on OG-pretreated compared to water-pretreated *bak1-5 bkk1-1* double mutant plants (Fig. 2b). This finding suggests that BAK1 and BKK1 are essential for the OG-induced reduction of aphid fecundity on Arabidopsis, consistent with their established role as positive regulators of plant immunity (Roux *et al*., 2011). Nonetheless, *bak1-5 bkk1-1* mutant plants were overall more resistant to aphids than Col-0 WT plants (Fig. 2b). This is consistent with ETI being activated in *bak1 bkk1* mutants in a light-and SA-dependent manner (Gao *et al*., 2017; Wu *et al*., 2020).

**Fig. 2.**
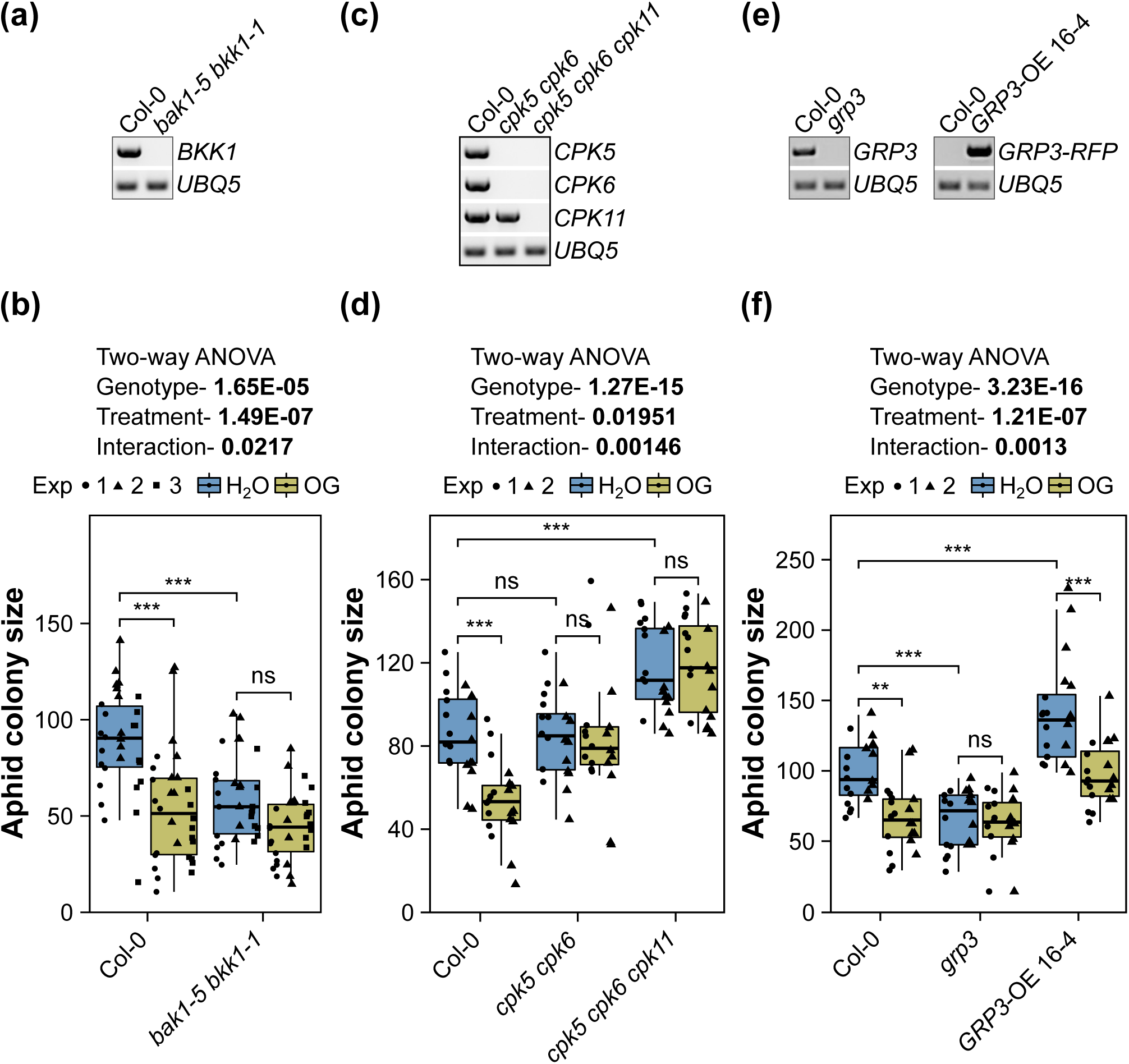
The oligogalacturonide (OG)-induced protection against Myzus persicae requires brassinosteroid-insensitive 1 (BRI1)-associated receptor kinase 1 (BAK1) and BAK1-like (BKK1), calcium-dependent protein kinases (CDPKs) CPK5 and CPK6, and glycine-rich protein 3 (GRP3). (a, c, e) Transcript analyses in bak1-5 bkk1-1, cpk5 cpk6, cpk5 cpk6 cpk11, and grp3 mutants, and in GRP3-OE transgenic plants, compared to Col-0 WT, using gene-specific primers listed in Supporting Information Table S1 and complementary DNA (cDNA) as a template. (b, d, f) Aphid single leaf fecundity assays. 4-weeks (w)-old plants of Arabidopsis thaliana Col-0 wild type (WT) and bak1-5 bkk1-1, cpk5 cpk6, cpk5 cpk6 cpk11, and grp3 mutants, and in GRP3-OE transgenic plants, as indicated, were infiltrated with H_2_O or 200 μg ml^-1^ OG. 72 h post infiltration, one 6-d-old asexually reproducing M. persicae adult female was caged on the infiltrated leaf. After 10 d, the number of individual aphids (adults + nymphs) within the cage was counted to obtain the aphid colony size. The y-axis shows the aphid colony size. The x-axis shows the plant genotype. Boxplots show the median, the 25^th^ and 75^th^ percentiles, the most extreme data points (whiskers’ extensions), and the observations as black-filled circles (experiment [exp] 1), triangles (exp 2), or squares (exp 3). n = 10 in each experiment. Asterisks indicate significant differences between samples as determined by two-way ANOVA with interaction (genotype:treatment) and post-hoc Tukey HSD test (**, P < 0.01; ***, P < 0.001; ns, not significant).

CDPKs have a role in OG-induced DTI downstream of BAK1, with CPK5, CPK6, and CPK11 playing a redundant and equal contribution (Gravino *et al*., 2015). The OG pretreatment did not reduce aphid fecundity on *cpk5 cpk6* double and *cpk5 cpk6 cpk11* triple mutants (Fig. 2c,d), whereas aphid fecundity was reduced on the OG-pretreated Col-0 WT plants (Fig. 2d). Moreover, the aphids produced more progeny on the *cpk5 cpk6 cpk11* triple mutant versus Col-0 WT (Fig. 2d). These data indicate that the three CPKs play a role in OG-induced resistance to aphids, with CPK11 having a more prominent role in mediating basal resistance of Arabidopsis to *M. persicae*.

GRP3 is a negative regulator of OG-induced DTI (Gramegna *et al*., 2016). Consequently, an Arabidopsis *grp3 null* mutant is more resistant to *Botrytis cinerea* (Gramegna *et al*., 2016). In our experiments, aphids produced less progeny on the *grp3 null* mutant (Fig. 2e) compared to Col-0 WT plants (Fig. 2f), indicating that the *grp3 null* mutant is also more resistant to *M. persicae*. Moreover, compared to Col-0 WT plants, the aphids produced more progeny on an Arabidopsis line that overexpressed *GRP3* as a fusion to *RFP* under the control of the *35S* promoter (*35S:GRP3::RFP* #16, Fig. 2e) (Gramegna *et al*., 2016) (Fig. 2f). The OG pretreatment did not affect aphid fecundity on the *grp3 null* mutant but reduced aphid fecundity on the *35S:GRP3::RFP* and Col-0 WT plants, as compared to water-pretreated plants (Fig. 2f). These data suggest that GPR3 positively affects the OG-induced reduction of aphid fecundity on Arabidopsis and acts as a susceptibility factor in the basal resistance of Arabidopsis to *M. persicae*.

Altogether, these results show that BAK1/BKK1, CPK5/CPK6, and GRP3 play an essential role in the OG-induced DTI that reduces the ability of *M. persicae* to colonise Arabidopsis plants.

### Early and late aphid feeding minimises *in vivo* accumulation of long OGs in Arabidopsis

Given the critical role of OGs in plant defences against *M. persicae*, it remains unclear how aphid feeding affects their accumulation. To address this, we quantified the effect of aphid feeding on the capacity of the plant to produce OGs. We measured OG abundance in leaf diffusates leaking from leaf strips; this method of wounding induces the production and release of OGs (Savatin *et al*., 2014b). Extracts were prepared from Arabidopsis Col-0 WT leaves previously infested for 6 h with aphids, confined to individual leaves using clip cages, compared to empty clip cages or untreated controls (Fig. 3a). This experimental setup was chosen based on evidence that pectin changes occur within 6 h of *M. persicae* infestation and that aphids settle and accept Arabidopsis as a host within 3-6 h of their initial probe (Silva-Sanzana *et al*., 2019).

**Fig. 3.**
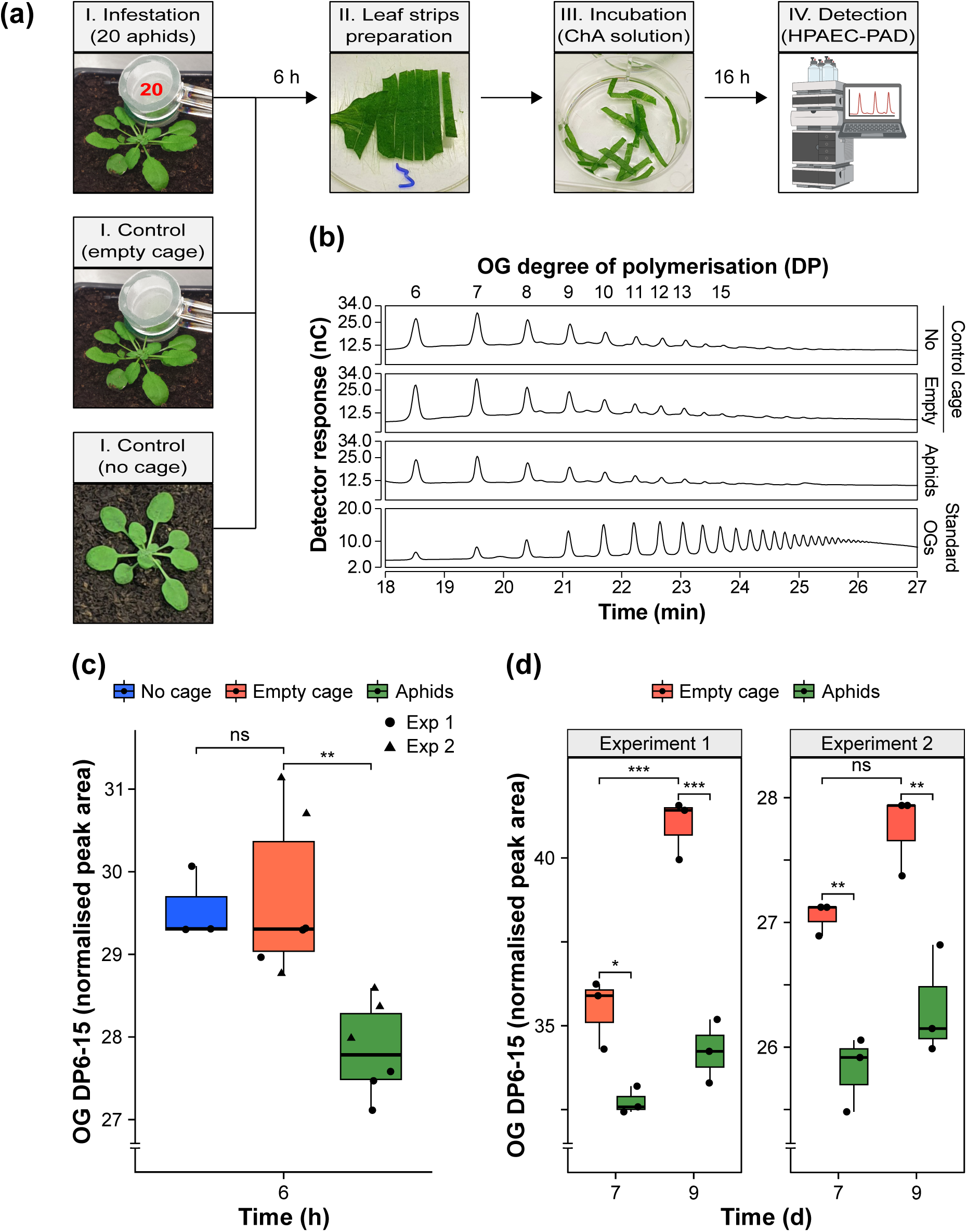
*In vivo* accumulation of long oligogalacturonides (OGs) is suppressed during *Myzus persicae* colonisation of Arabidopsis. (a) Experimental design. Leaves from 4-week-old Arabidopsis Col-0 wild-type (WT) plants were caged with twenty 6-d-old asexually reproducing *M. persicae* adult females, without aphids (empty cage), or not caged, as a control (panels I). After 6 h, leaves were excised and, after removing all aphids, sliced into strips (panel II). Leaf strips were incubated with strong chelating agents supplemented with sodium sulphite (ChA solution) for 16 h (panel III). The leaf diffusates in incubation medium were analysed by high-performance anion-exchange chromatography (HPAEC) with a pulsed amperometric detector (PAD) (panel IV). The illustration in panel IV was created with BioRender.com (Agreement number: EI25PBPRUZ). (b) Chromatographic analyses show the profile of OGs across samples. The numbers on top of the graph indicate the degree of polymerisation (DP) of different OG oligomers and refer to the corresponding peaks below. The x-axis shows the retention time in min. The y-axis shows the detector response in nanocoulombs (nC). For each sample, the result of one representative replicate out of three performed is shown. The bottom panel shows the profile of 2.5 μg of standard OGs enriched in DP10-15. (c, d) Total content of OGs (DP6-15) in control samples and aphid-infested leaves for 6 h (c) or 7 and 9 d (d). The x-axis shows the time of infestation in h or d. The y-axis shows the peak area (nC min) of total OGs from DP6 to DP15, normalised using the compositional data normalisation (CoDA) method. Boxplots show the median, the 25^th^ and 75^th^ percentiles, the most extreme data points (whiskers’ extensions), and the observations as black-filled circles. Two independent experiments are shown. n = 3 in each experiment. Asterisks indicate significant differences between samples as determined by one-way ANOVA with post-hoc Tukey HSD test (*, *P* < 0.05; **, *P* < 0.01; ***, *P* < 0.001).

HPAEC-PAD analysis of leaf diffusates identified peaks in all samples that eluted with retention times comparable to those of an OG solution enriched in long OGs with DP10-15 (Benedetti *et al*., 2017) (Fig. 3b), which most effectively induce plant defences (Mathieu *et al*., 1991; Ferrari *et al*., 2013). However, quantification of the area under the peaks and data normalisation showed that total OG levels with DP6-15 were lower in diffusates collected from aphid-infested leaves compared to clip cage controls, with no differences observed between diffusates collected from leaves treated with or without empty clip cages (Fig. 3c; Supporting Information Fig. S3a).

To account for the possible impact of late aphid feeding on OG release, OG abundance was also assessed after 7 and 9 d of aphid infestation, corresponding to the time points of aphid fecundity assays (Fig. 1b,f). In these experiments, Arabidopsis Col-0 WT leaves were infested with clip cages containing two adult aphids or exposed to empty clip cages as a control (Supporting Information Fig. S3b). However, despite an average of 30-38 aphids produced per female by these time points (Supporting Information Fig. S3c), total OG levels with DP6-15 were consistently lower in diffusates collected from aphid-infested leaves compared to controls after both 7 and 9 d (Fig. 3d; Supporting Information Fig. S3d,e). In one experiment, we identified OGs with DP3, which exhibit a weaker immune activity (Davidsson *et al*., 2017), and observed that their levels were also reduced in diffusates collected from aphid-infested leaves compared to controls after both 7 and 9 d (Supporting Information Fig. S3e). Notably, we detected higher levels of OGs in control leaves sampled at 9 days compared to those sampled at 7 days (Fig. 3d), as previously observed in plants at different developmental stages and potentially due to cell wall remodelling associated with plant growth and development (Pontiggia *et al*., 2015; Pontiggia *et al*., 2020). Shifted peaks potentially corresponding to oxidised OGs that are known not to activate DTI (Benedetti *et al*., 2018) were not detectable in any of our experiments (Supporting Information Fig. S4a,b,c,d). In all our experiments, some peaks of unknown identity were reduced, while others were unchanged or increased in aphid-infested leaves (Supporting Information Fig. S3a,e).

In summary, defence-inducing OGs are released in Arabidopsis aphid-infested leaves upon wounding but in lower amounts than in non-infested leaves.

### The aphid effector Mp10 suppresses the OG-triggered ROS burst, defence gene induction, and resistance against *M. persicae*

The aphid effector Mp10 is delivered into the cytoplasm of plant mesophyll cells during the early probing phase of aphid feeding (Mugford *et al*., 2016) and has been shown to suppress plant PTI responses (Bos *et al*., 2010; Drurey *et al*., 2019; Gravino *et al*., 2024). Here, we investigated if Mp10 also suppresses DTI responses. In Arabidopsis Col-0 WT leaves, OGs induced typical DTI responses, including a peak of ROS within 10 m following elicitation returning to near-basal levels after 30 m (Fig. 4a), that contributed to the total ROS productions (Fig. 4b), in agreement with previous data (Gravino *et al*., 2015; Gravino *et al*., 2017). Moreover, treatment with OGs induced the expression of the defence-related genes *FRK1*, *CYP81F2*, *PAD3*, and *PAD4* (Fig. 4c), as previously reported (Denoux *et al*., 2008; Gravino *et al*., 2015; Gravino *et al*., 2017). To investigate whether Mp10 suppresses OG-induced ROS and defence genes, we generated stable transgenic plants that express Mp10 under the control of a DEX-inducible promoter in the Arabidopsis Col-0 background. Two independent homozygous lines (#7-5 and #9-5) that showed *Mp10* expression upon treatment with DEX (solubilised in DMSO) and not with DMSO alone were selected (Supporting Information Fig. S5a,b). ROS kinetics over time and total ROS productions, as well as the expression of the defence genes *FRK1*, *CYP81F2*, *PAD3*, and *PAD4* in response to OGs, were reduced in Mp10 lines pretreated with 1.25 μM DEX compared to Col-0 plants pretreated with 1.25 μM DEX (Fig. 4a,b,c), indicating that Mp10 suppresses the ROS burst and defence genes induced by OGs.

**Fig. 4.**
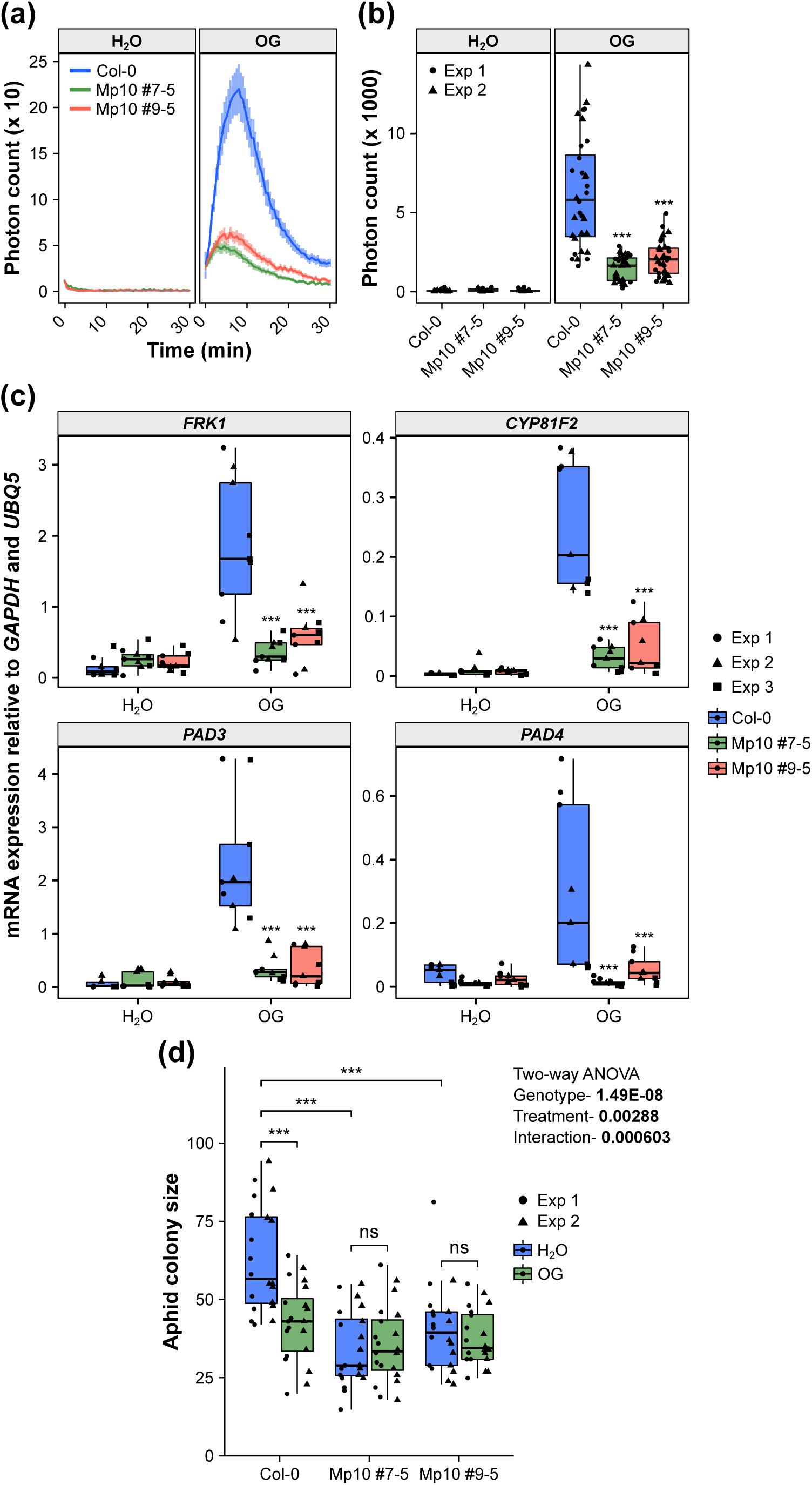
The aphid effector Mp10 suppresses the oligogalacturonide (OG)-triggered immunity. In (a, b, c, d), 3- to 4-week (w)-old plants of *Arabidopsis thaliana* Col-0 wild type (WT) and dexamethasone (DEX)-inducible Mp10 transgenic lines (#7-5 and #9-5) were sprayed with 1.25 μM DEX 3 d before treatment with 200 μg ml^-1^ OG and H_2_O as a control. (a) ROS production measured in leaf discs over a period of 30 min after elicitor treatment. Solid line, mean; shaded band, SEM. *n* = 16 and 32 leaf discs treated with H_2_O and OGs, respectively, from two independent experiments. The x-axis shows the time in min after elicitor treatment. The y-axis shows the photon count. (b) Total ROS production over a period of 30 min after elicitor treatment. *n* = 8 and 16 leaf discs treated with H_2_O and OGs, respectively, in each experiment. The y-axis shows the total photon count. The x- axis shows the plant genotype. (c) Transcript levels of *FLG22-INDUCED RECEPTOR- LIKE KINASE 1* (*FRK1*), *CYTOCHROME P450, FAMILY 81, SUBFAMILY F, POLYPEPTIDE 2* (*CYP81F2*), *PHYTOALEXIN DEFICIENT 3* (*PAD3*), and *PHYTOALEXIN DEFICIENT 4* (*PAD4*) relative to *GLYCERALDEHYDE 3-PHOSPHATE DEHYDROGENASE* (*GAPDH*) and *UBIQUITIN 5* (*UBQ5*) (y-axis) measured 2 h after elicitor infiltration. *n* = 3 in each experiment. (d) Aphid single leaf fecundity assay. 72 h post-elicitor infiltration, one 6-d-old asexually reproducing *M. persicae* adult female was caged on the infiltrated leaf. After 10 d, the number of individual aphids (adults + nymphs) within the cage was counted to obtain the aphid colony size. The y-axis shows the aphid colony size. The x-axis shows the plant genotype. In (b, c, d), boxplots show the median, the 25^th^ and 75^th^ percentiles, the most extreme data points (whiskers’ extensions), and the observations as black-filled circles in experiment (exp) 1, triangles in exp 2, or squares in exp 3. *n* = 10 in each experiment. Asterisks indicate significant differences between samples as determined by one-way ANOVA or two-way ANOVA with interaction (genotype:treatment) and post-hoc Tukey HSD test, as indicated (***, *P* < 0.001; ns, not significant).

To further explore the specificity of Mp10 suppression of elicitor-induced plant immunity, we assessed whether Mp10 inhibits ROS burst responses triggered by bacterial flg22 and fungal chitin, which activate BAK1-dependent and BAK1-independent immune signalling, respectively (Shan *et al*., 2008). In Mp10-expressing lines pretreated with 1.25 µM DEX, ROS bursts induced by flg22 were significantly reduced compared to Col-0 plants similarly pretreated with DEX (Supporting Information Fig. S6a). In contrast, chitin-induced ROS bursts were unaffected in Mp10 lines (Supporting Information Fig. S6b), aligning with previous findings (Bos *et al*., 2010). These results confirm that Mp10 specifically suppresses BAK1-dependent immune-signalling pathways while leaving BAK1-independent pathways intact.

To test if the suppression of the OG-triggered DTI by Mp10 favours aphid colonization, we measured aphid fecundity in Arabidopsis Col-0 WT and Mp10-expressing plants pretreated with 1.25 μM DEX prior to treatment with or without OGs. In contrast to Col-0 WT plants, aphid fecundity was not reduced on OG-treated compared to water-treated Mp10 lines (#7-5 and #9-5) (Fig. 4d), suggesting that Mp10 suppresses the OG-induced reduction of aphid fecundity on Arabidopsis. Nonetheless, Mp10-expressing lines were overall more resistant to aphids than Col-0 WT plants (Fig. 4d), in agreement with previous data (Bos *et al*., 2010).

### Mp10 induces immune responses that are differentially dependent on EDS1, ADR1, NRG1, and salicylic acid

Since Mp10 induces immunity in a salicylic acid glucosyltransferase 1 (SGT1)-, RCSP-, and EDS1-dependent manner (Bos *et al*., 2010; Rao *et al*., 2024), we further investigated if ACTIVATED DISEASE RESISTANCE 1 (ADR1), N REQUIREMENT GENE 1 (NRG1), and salicylic acid (SA) are involved in this pathway. Transient production of Mp10 *via* agroinfiltration in *N. benthamiana* WT leaves induced localised chlorosis in the infiltrated areas observed as early as 3 days post infiltration (dpi) (Fig. 5a,b) (Bos *et al*., 2010; Drurey, 2015). This was followed by systemic chlorosis in uninfiltrated leaves and an overall dwarf phenotype at approximately 14 dpi (Fig. 5c) (Bos *et al*., 2010; Drurey, 2015; Rao *et al*., 2024). The local chlorosis was observed in the *N. benthamiana nrg1-1* and *nrg1-2* mutants and partially in NahG plants (Fig. 5a,b) – NahG encodes a bacterial salicylate hydroxylase that destroys SA (Abreu & Munne-Bosch, 2009). However, it was absent in *eds1* and *adr1 nrg1* mutants (Fig. 5a,b). In contrast, systemic chlorosis and dwarfism were not observed in *N. benthamiana nrg1-1*, *nrg1-2*, *eds1* mutants, or NahG plants (Fig. 5c,d).

**Fig. 5.**
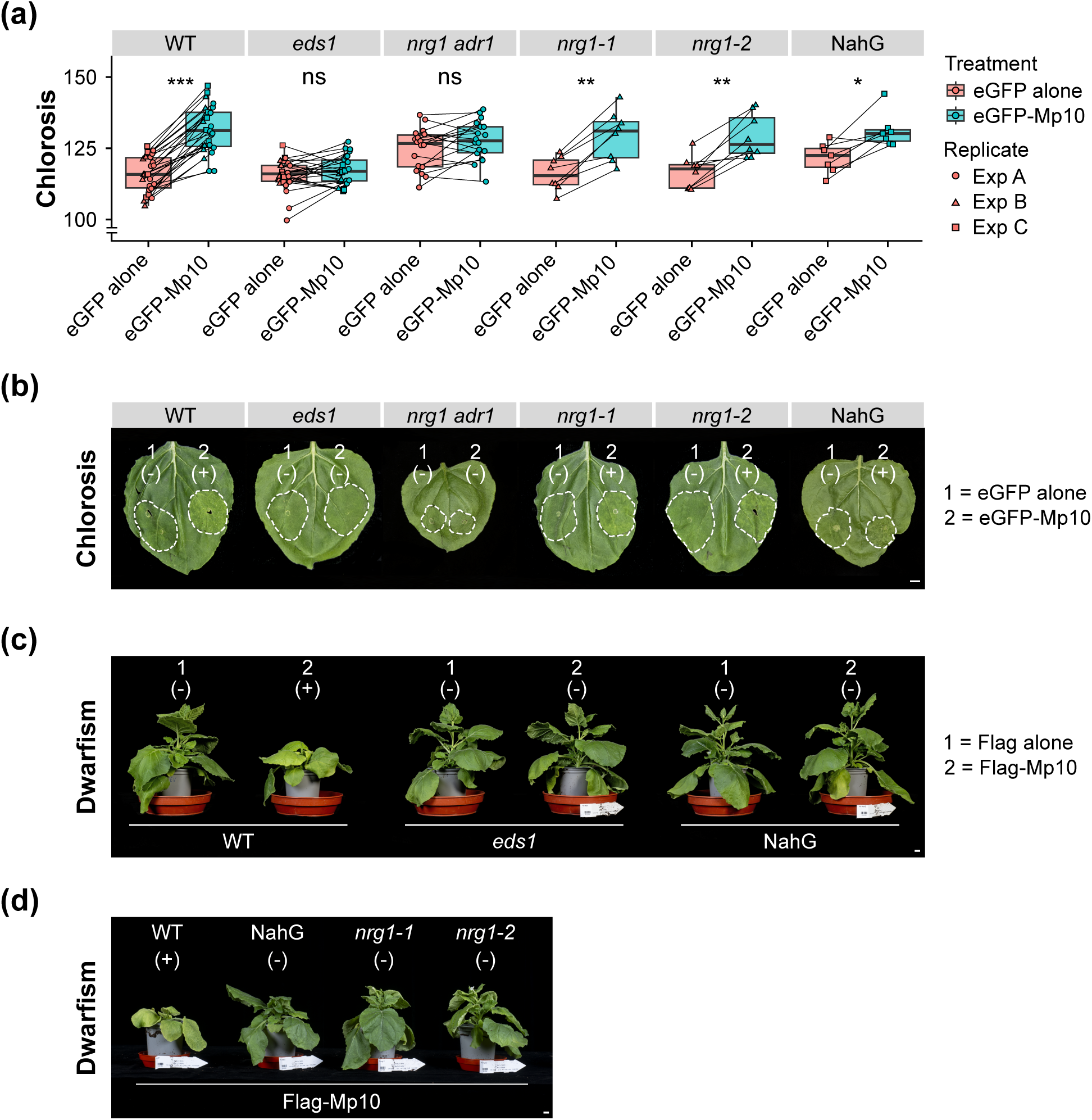
The aphid effector Mp10 promotes effector-triggered immunity (ETI)-like responses that are differentially dependent on EDS1, ADR1, NRG1, and salicylic acid. (a) Chlorosis measurements in *Nicotiana benthamiana* leaves from WT and the indicated mutants agroinfiltrated with either eGFP alone or eGFP-Mp10 (x-axis). Chlorosis, defined as the yellowing of normally green leaf tissue, is quantified on the y-axis as the average of the red and green colour components in the agroinfiltrated areas. Boxplots show the median, the 25^th^ and 75^th^ percentiles, the most extreme data points (whiskers’ extensions), and the observations as circles in experiment (exp) 1, triangles in exp 2, or squares in exp 3. *n* ≥ 7 in each experiment. For each genotype, asterisks indicate significant differences between treatments as determined by two-way ANOVA with interaction (genotype:treatment) and post-hoc Tukey HSD test, as indicated (***, *P* < 0.001; **, *P* < 0.01; *, *P* < 0.05; ns, not significant). (b) Representative pictures of *N. benthamiana* leaf patches showing chlorosis (+) or not (-) after agroinfiltration with either eGFP alone (1) or eGFP-Mp10 (2) on each side of one leaf from WT and the indicated mutants. Pictures were taken 7 days post- infiltration (dpi). (c, d) *N. benthamiana* plants of WT and indicated mutants showing systemic chlorosis and dwarfism (+) or not (-) after agroinfiltration with either Flag alone (1) or Flag-Mp10 (2), as indicated, on one side of one leaf. Pictures were taken at 14 dpi. (b, c, d) Scale bars = 1 cm.

Similarly, in Arabidopsis Mp10 transgenic lines, we observed chlorosis/senescence, dwarfism, and intracellular H_2_O_2_ accumulation, as detected by DAB staining. These symptoms were more pronounced in the Mp10 line #7-5 compared to line #9-5, which expresses lower levels of *Mp10* (Supporting Information Fig. S5a,b) and only exhibited symptoms under stress conditions (e.g., continuous darkness). Notably, these phenotypes were absent in Arabidopsis *eds1-2* x Mp10 lines, as well as in WT and *eds1-2* mutant plants (Supporting Information Fig. S7a,b,c,d,e,f,g,h).

These findings indicate that Mp10 triggers ETI-like responses that are dependent on EDS1, ADR1, NRG1, and SA.

### Mp10 modulates EDS1-associated processes as part of the DTI response while also activating ETI

Since Mp10 triggers immunity in Arabidopsis in an EDS1-dependent manner, potentially masking other Mp10-related activities, we further examined its impact on OG-induced immunity to aphids in the absence of EDS1. Aphids showed reduced fecundity on OG-treated WT plants compared to water-treated WT plants, but their performance remained similar on water- and OG-treated Arabidopsis *eds1-2* plants (Fig. 6a,b), indicating that OG- induced DTI depends on EDS1. This finding is consistent with previous reports demonstrating a role for EDS1 in pattern recognition receptor signalling in Arabidopsis (Pruitt *et al*., 2021). Here, we extend this role to OG-induced signalling, suggesting that EDS1 is also required for the defence response triggered by OGs.

**Fig. 6.**
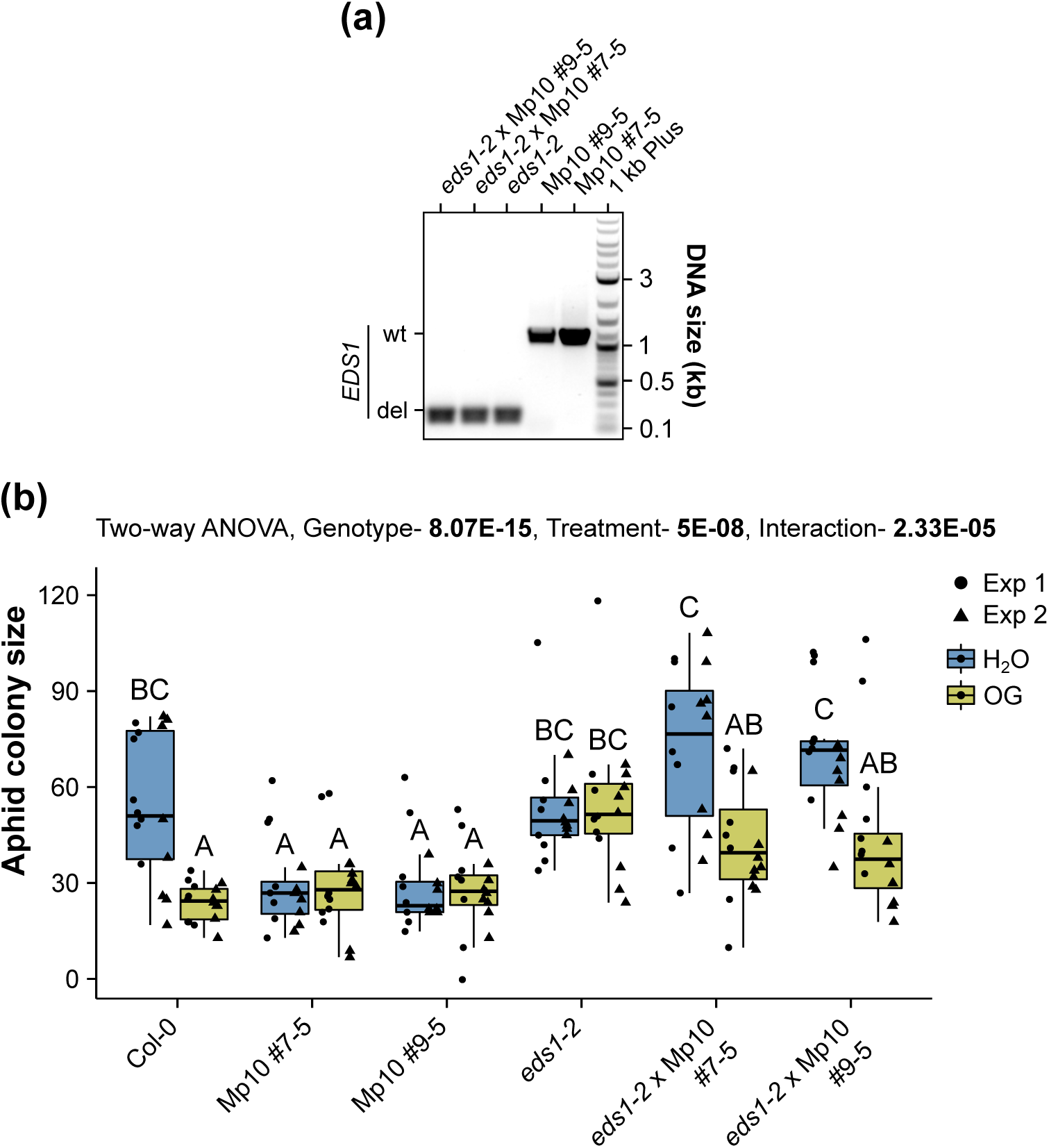
The aphid effector Mp10 promotes aphid resistance and suppresses the oligogalacturonide (OG)-induced protection against *Myzus persicae* in an *EDS1*- dependent manner. (a) Analysis of the *EDS1* deletion in the *eds1-2* mutant, *eds1-2* x Mp10 #7-5, and *eds1-2* x Mp10 #9-5 crosses, compared to Mp10 transgenic lines (#7-5 and #9-5), using *EDS1*-specific primers, flanking the deleted region, listed in Supporting Information Table S1, and genomic DNA as a template. (b) Aphid fecundity assay. 3- to 4- week (w)-old plants of *Arabidopsis thaliana* Col-0 wild type (WT) and dexamethasone (DEX)-inducible Mp10 transgenic lines (#7-5 and #9-5), *eds1-2* mutant, *eds1-2* x Mp10 #7- 5, and *eds1-2* x Mp10 #9-5 crosses, as indicated, were sprayed with 1.25 μM DEX and, 3 d later, infiltrated with H_2_O and 200 μg ml^-1^ OG. 72 h post infiltration, one 6-d-old asexually reproducing *M. persicae* adult female was caged on the infiltrated leaf. After 10 d, the number of individual aphids (adults + nymphs) within the cage was counted to obtain the aphid colony size. The y-axis shows the aphid colony size. The x-axis shows the plant genotype. Boxplots show the median, the 25^th^ and 75^th^ percentiles, the most extreme data points (whiskers’ extensions), and the observations as black-filled circles in experiment (exp) 1 or triangles in exp 2. n = 8 samples in each experiment. Different letters indicate significant differences between samples as determined by two-way ANOVA with interaction (genotype:treatment) and post-hoc Tukey HSD test. See Table S3 for P-values.

Aphid performances on the water- and OG-treated Mp10 lines were similar and comparable to those of OG-treated WT plants (Fig. 6b), in agreement with previous data (Fig. 4d). However, aphids did less well on OG-treated than on water-treated *eds1-2* x Mp10 lines (Fig. 6b), indicating that the OG-mediated resistance response to aphids was present in the *eds1-2* x Mp10 lines, unlike the *eds1-2* lines. Moreover, aphids produced more progeny on the *eds1-2* x Mp10 lines compared to Mp10 lines (Fig. 6b), consistent with our finding that the EDS1 node is required for Mp10-induced ETI (Fig. 5). Aphids produced more progeny on water and OG-treated *eds1-2* x Mp10 lines versus WT plants, but the differences were not statistically significant.

Overall, these data indicate that Mp10 promotes aphid fecundity in the absence of EDS1 and that EDS1 is required for OG-induced immunity against aphids. However, Mp10 compensates for the loss of EDS1, restoring plant responsiveness to OGs. Therefore, Mp10 operates within the EDS1 pathway, likely modulating EDS1-associated processes as part of the DTI response while also activating ETI.

### EDS1 is crucial for Mp10-mediated PRR destabilisation and PTI suppression

Our finding that Mp10 may act in the EDS1 node instigated further investigations of the involvement of EDS1 in Mp10 immune-suppressing activities. Since it is unclear which PRRs perceive OGs (Herold *et al*., 2024), we used the well-characterised flg22/FLS2 system, as it was previously found that flg22 treatment induces the expression of immune genes effective against aphids (Kettles *et al*., 2013; Prince *et al*., 2014a) and Mp10 affects the stability of the flg22 receptor FLS2 (Gravino *et al*., 2024). Mp10 suppressed the flg22- induced ROS burst in WT *N. benthamiana* leaves as observed previously (Bos et al., 2010), whereas this Mp10-mediated suppression activity was not observed in *N. benthamiana eds1* plants (Fig. 7a). EDS1 is not known to impact flg22-elicited ROS production (Fig. 7a) (Zönnchen *et al*., 2022). Moreover, Mp10 destabilised FLS2 in WT *N. benthamiana*, consistent with previous findings (Gravino *et al*., 2024), but not in *N. benthamiana eds1* plants (Fig. 7b). These data indicate that EDS1 is required for the Mp10-mediated ROS suppression and PRR destabilisation activities.

**Fig. 7.**
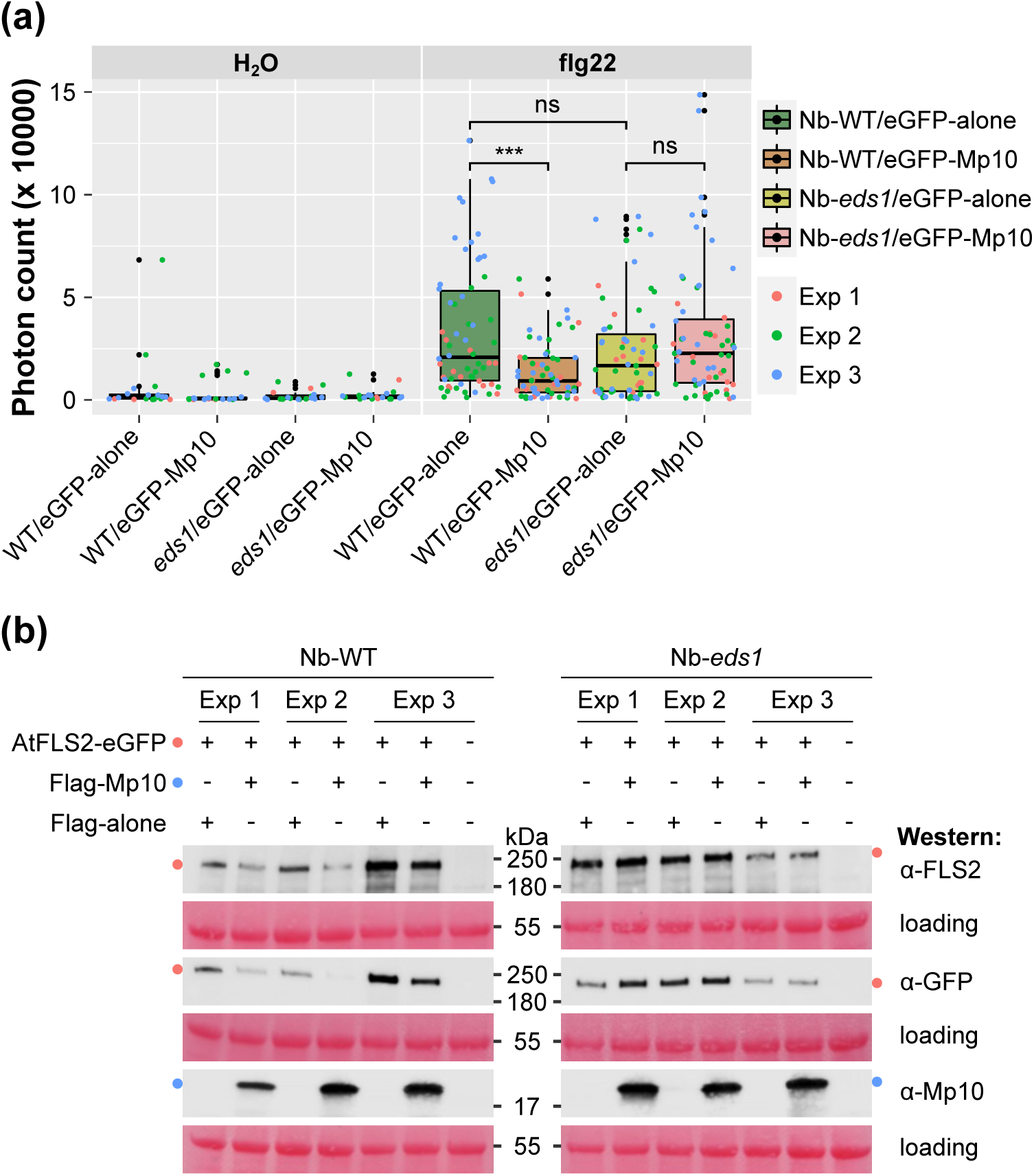
EDS1 is essential for Mp10-mediated suppression of the reactive oxygen species (ROS) burst induced by bacterial flg22 and the destabilisation of FLS2. (a) Total ROS production over 1 h following treatment with 100 nM flg22 or H₂O (control) in *Nicotiana benthamiana* WT or *eds1* mutant leaf discs agroinfiltrated with eGFP or eGFP-Mp10. Y- axis: total photon count; X-axis: plant genotype/construct. Boxplots display the median, 25^th^ and 75^th^ percentiles, whiskers indicating extreme data points, and observations as red (exp1), green (exp2), or blue (exp3) circles. n ≥ 3, with 15 leaf discs per treatment per experiment. Asterisks indicate significant differences (one-way ANOVA, Tukey HSD; ***, P < 0.001; ns, not significant). (b) Western blot analysis of *N. benthamiana* WT or *eds1* mutant leaf discs co-agroinfiltrated with Arabidopsis FLS2-eGFP and either Flag or Flag- Mp10. FLS2-eGFP protein stabilisation in the presence/absence of Mp10 was detected using FLS2 (top), eGFP (middle), and Mp10 (bottom) antibodies. MW markers are shown on the right (WT) or left (*eds1*), with expected bands marked by coloured dots. Loading control: Ponceau S staining.

## DISCUSSION

This study sheds light on the intricate dynamics of DTI activation with a specific focus on the roles of OGs as DAMPs and DTI suppression by the aphid saliva chemosensory protein effector Mp10 (Fig. 8). We found that OG-induced DTI effectively limits *M. persicae* colonisation on Arabidopsis and that the key immune components BAK1/BKK1, CPK5/CPK6, GRP3, and EDS1 are pivotal for this DTI response. However, aphids employ diverse strategies to counteract these defences. Firstly, OG production upon wounding is reduced in plants exposed to aphid feeding, indicating that aphids may suppress DAMP release, thereby enhancing their performance. Moreover, the aphid effector Mp10, which is secreted into the cytoplasm of plant cells during the early stages of aphid feeding (Mugford *et al*., 2016), suppresses the plant sensitivity to OGs. However, Mp10 also induces ETI, including local and systemic responses, in an EDS1-dependent manner. Remarkably, we found that EDS1 is required for OG-induced DTI and Mp10-mediated ROS suppression and PRR destabilisation activities, whereas Mp10 restores OG-induced DTI in the absence of EDS1.

**Fig. 8.**
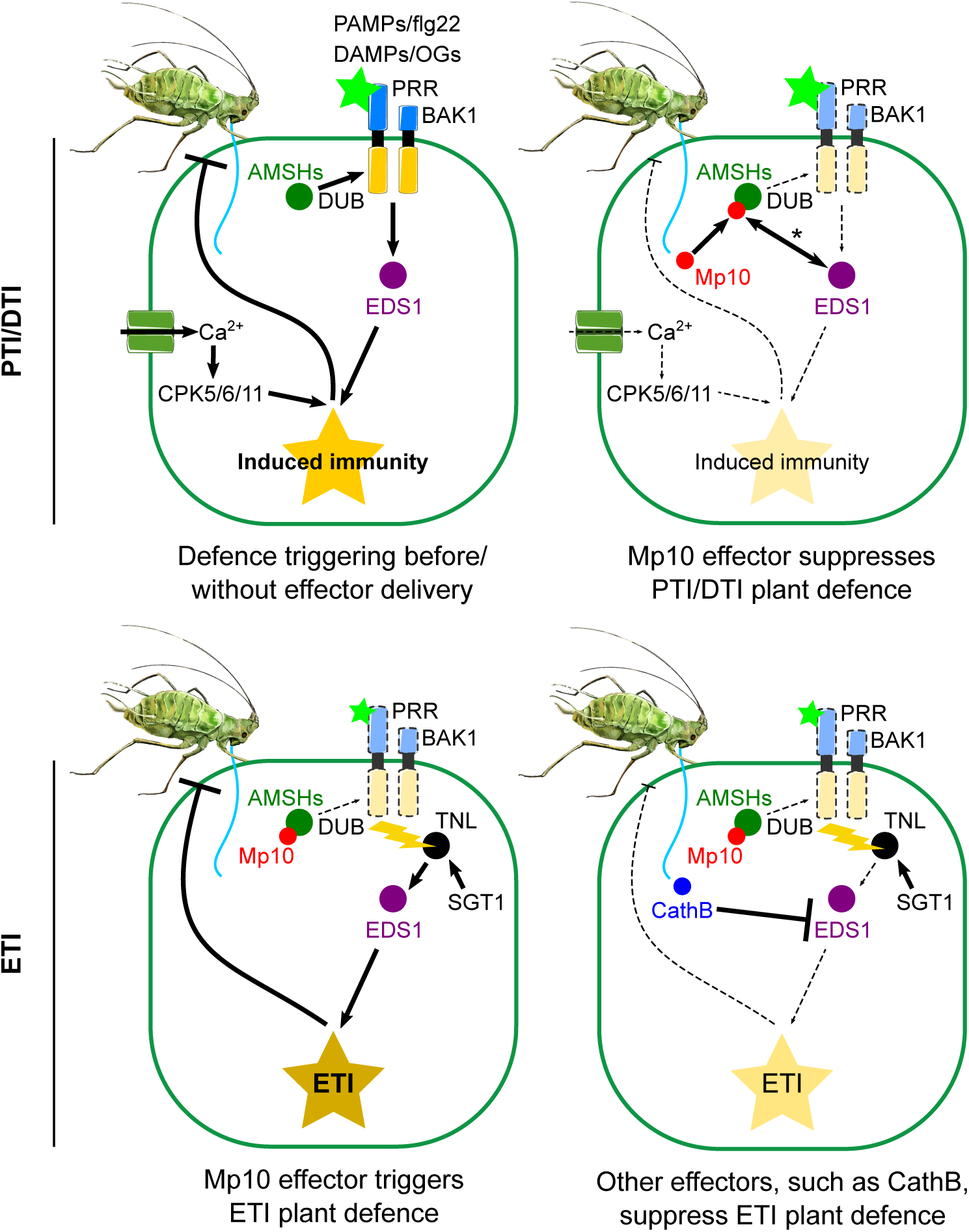
Model: Activation and suppression of defences in plant-aphid interactions. <u>Top Left</u> <u>Panel</u>: Aphid stylet penetration of the cell wall releases oligogalacturonides (OGs), which induce PAMP/DAMP-triggered immunity (PTI/DTI) in a process dependent on BAK1, EDS1, and CPK5/6/11 (data herein). The stabilisation of pattern recognition receptors (PRRs) and coreceptors at the plasma membrane may rely on the deubiquitination (DUB) activity of AMSHs (Gravino *et al*., 2024). <u>Top Right Panel</u>: The Mp10 effector, introduced by aphids into the cell cytoplasm (Mugford *et al*., 2016), suppresses OG-induced ROS (data herein) and flg22-induced PTI (Bos *et al*., 2010). Mp10 targeting of plant AMSHs is implicated in these processes (Gravino *et al*., 2024). EDS1 is essential for Mp10-mediated ROS suppression and PRR destabilisation (data herein), acting through an unidentified mechanism (denoted by the double-sided arrow with an asterisk). <u>Bottom Left Panel</u>: Effector-triggered immunity (ETI) is activated through EDS1, either directly or indirectly, upon recognition of Mp10 and/or its activities by a TNL (Gravino *et al*., 2024; Rao *et al*., 2024). SGT1 is also required for TNL/ETI activation (Bos *et al*., 2010). <u>Bottom Right Panel</u>: Aphids secrete additional effectors, such as cathepsin B proteins (e.g., CathB6), which target EDS1 to suppress ETI (Liu *et al*., 2024). The model was generated using icons with attribution and licensing details in Table S3.

OG-induced DTI against *M. persicae* depends on BAK1/BKK1, GRP3, EDS1, and CPK5/6, while CPK11 promotes plant immunity to aphids. These kinases are members of the CDPK subgroup I (Cheng *et al*., 2002) and are known regulators of defence responses downstream of various PAMPs/DAMPs, contributing to resistance against pathogens such as *P. syringae* and *B. cinerea* (Boudsocq *et al*., 2010; Dubiella *et al*., 2013; Ma *et al*., 2013; Gravino *et al*., 2015). Activated by calcium, CPK5/6/11 phosphorylate RBOHD *via* interaction with the lectin receptor-like kinase LecRK-IX.2, triggering PTI/DTI responses including ROS production and ethylene biosynthesis (Boudsocq *et al*., 2010; Kadota *et al*., 2014; Luo *et al*., 2014; Gravino *et al*., 2015; Luo *et al*., 2017). Aphid performance is unchanged on *cpk5 cpk6* double mutants but improves on *cpk5 cpk6 cpk11* triple mutants, and therefore ROS and ethylene production mediated by CPK5/6/11 may be crucial for aphid resistance. Additionally, BAK1/BKK1 and EDS1 function in related pathways, acting upstream and downstream of calcium elevations, respectively (Ranf *et al*., 2011; Roux *et al*., 2011; Pruitt *et al*., 2021; Tian *et al*., 2021). Consistent with this, *M. persicae* induces calcium elevations in wild-type plants that require BAK1 and performs better on mutants impaired in ROS production (*rbohd*) and ethylene signalling (*ein2*) (Moloi & van der Westhuizen, 2006; Miller *et al*., 2009; Kettles *et al*., 2013; Vincent *et al*., 2017). Together, these findings highlight the central role of BAK1/BKK1, GRP3, EDS1, and CPK5/6/11 in coordinating plant resistance to aphids.

We found that OG release is minimised in plants exposed to aphids. One possible explanation is that pectin becomes less susceptible to enzymatic degradation by either aphid salivary enzymes or plant enzymes activated upon wounding. Evidence suggests that plant PME activity increases during aphid feeding on Arabidopsis, leading to blockwise de-methylesterification and calcium-mediated cross-linking of pectin into “egg-box” structures (Silva-Sanzana *et al*., 2019). These structural modifications likely hinder the accessibility of pectin-degrading enzymes, promoting pectin stiffening and reducing OG release (Liners *et al*., 1989; Hocq *et al*., 2017; Wormit & Usadel, 2018). Another consideration is that OGs that are released by epidermal and mesophyll cells may be absorbed by the gel-like sheath saliva that surrounds the stylets of aphids within the apoplast. Such absorption may prevent binding of OGs to cell-surface receptors and reduce the detection of them using methods herein. Nevertheless, our findings suggests that aphids may actively minimise the release of immune-activating OGs to enhance their performance on host plants.

OG-induced immunity shares several characteristics with flg22. Mp10 suppresses OG- and flg22-induced immunity (Bos *et al*., 2010; Drurey *et al*., 2019; Gravino *et al*., 2024), which requires BAK1/BKK1, CPK5/CPK6/CPK11, GRP3, and EDS1 (Boudsocq *et al*., 2010; Roux *et al*., 2011; Gramegna *et al*., 2016; Pruitt *et al*., 2021; Tian *et al*., 2021). This overlap further suggests that OGs and flg22 share similar signal transduction elements (Gravino *et al*., 2017). In line with this, flg22 treatment also induced the expression of immune responses that reduce aphid performance (Prince *et al*., 2014a). Nonetheless, Mp10 suppression activity is specific because we found that this effector does not suppress chitin-induced PTI, in line with previous results (Bos *et al*., 2010). Interestingly, while both OG- and flg22-induced responses require BAK1 (Chinchilla *et al*., 2007; Heese *et al*., 2007; Gravino *et al*., 2017), chitin-induced responses are BAK1-independent (Shan *et al*., 2008). This distinction suggests that Mp10 specificity may involve BAK1-dependent pathways.

The aphid effector Mp10 exhibits dual roles in plant immunity, suppressing and inducing responses, a characteristic shared with pathogen effectors like AvrPto, AvrPtoB, and HopB1. The latter three effectors act by modulating FLS2 and other cell-surface receptor kinases (RKs). AvrPto inhibits their kinase activity (Xiang *et al*., 2008), while AvrPtoB and HopB1 promote the degradation of these receptors *via* ubiquitination or proteolytic cleavage (Goehre *et al*., 2008; Gimenez-Ibanez *et al*., 2009; Li *et al*., 2016), and the activities of these effectors are detected by intracellular immune receptors, leading to ETI (Mathieu *et al*., 2014; Schulze *et al*., 2022). Similarly, Mp10 destabilises immune-related RKs, including FLS2 and FER, in a process that involves interaction of Mp10 with AMSH proteins—key deubiquitinating enzymes essential for protein trafficking—thereby rendering plants insensitive to PAMPs (Gravino *et al*., 2024). Mp10 may also target putative OG receptors. However, the identity of these receptors remains elusive, as studies using WAK quintuple loss-of-function mutants showed that WAKs are entirely dispensable for the plant response to OGs (Herold *et al*., 2024) or that their absence might be compensated by redundant WAK-like or analogous receptors (Blaschek, 2024), leaving this hypothesis for future investigation. Given that depletion of RKs, including BAK1 and BKK1, from the plasma membrane is known to activate ETI *via* the EDS1 pathway (Gao *et al*., 2017; Wu *et al*., 2020; Schulze *et al*., 2022), our finding that Mp10 simultaneously suppresses PTI and triggers ETI is not unexpected. This is further supported by our observation that aphids exhibit reduced performance on Arabidopsis *bak1-5 bkk1* plants.

We found that Mp10-induced immunity exhibits differential dependency on EDS1, ADR1, NRG1, and SA. This includes a local transient expression of Mp10 inducing a weak response, characterised by chlorosis confined to the infiltration area, which is dependent on EDS1 and ADR1. This response does not require NRG1, which induces a strong immune activation, such as cell death (Lapin *et al*., 2019), in line with the absence of a clear cell death response to Mp10 in this and earlier studies (Bos *et al*., 2010; Rao *et al*., 2024). Additionally, the dependence of the local response on SA is consistent with the established role of EDS1 in mediating both SA-dependent and SA-independent immune pathways (Bartsch *et al*., 2006; Straus *et al*., 2010; Cui *et al*., 2017).

Mp10 triggers also a systemic effect in plants, as evidenced by a dwarfing phenotype of plants, and this phenotype is dependent on NRG1, as well as EDS1 and SA. Since EDS1 promotes ETI downstream of TIR-NBS-LRR (TNL) immune receptors (Aarts *et al*., 1998; Feys *et al*., 2001; Sun *et al*., 2021), our results suggest that Mp10 itself and/or its activities of perturbing cell-surface receptor kinases (Gravino *et al*., 2024) may be recognised/guarded by TNL immune receptors (Greenwood & Williams, 2022). Mp10 is a chemosensory protein that is present in other sap-feeding insects, including whiteflies, leafhoppers, and planthoppers (Drurey *et al*., 2019; Gravino *et al*., 2024). Recent evidence suggests that these CSPs are recognised by the *N. benthamiana* TNL-CJID (C-terminal jelly roll/Ig-like domain) protein Recognition of CSPs (RCSP), which also requires EDS1 for inducing chlorosis and dwarfing responses (Rao *et al*., 2024). More research is needed to assess if this receptor is involved in *M. persicae* Mp10-induced chlorosis and dwarfing responses reported herein.

Our results reveal that EDS1 is crucial for Mp10-mediated suppression of PTI and DTI. EDS1 is a conserved immune regulator essential for ETI and its crosstalk with PTI, linking the EDS1-PAD4-ADR1 complex to PRR complexes *via* SOBIR1 for downstream PTI signalling (Pruitt *et al*., 2021; Tian *et al*., 2021). It also monitors receptor kinase homeostasis involved in PTI and activates ETI when perturbations occur (Schulze *et al*., 2022; Yang *et al*., 2022; Yu *et al*., 2023). We found that Mp10 suppresses flg22-triggered PTI and destabilises FLS2, consistent with previous reports (Gravino *et al*., 2024). However, in the absence of EDS1, Mp10 no longer affects FLS2 stability or downstream signalling. Notably, OG-mediated DTI depends on EDS1 and is suppressed by Mp10. In Arabidopsis *eds1-2* x Mp10 cross lines, OG-mediated DTI immunity to aphids is restored, whereas it is absent in *eds1-2*, indicating that Mp10 compensates for the DTI response in the *eds1-2* mutant background. This suggests a complex interplay between Mp10 and EDS1 in regulating OG-inducible DTI. The mechanism by which EDS1 contributes to Mp10 suppressive activity is unclear. It is possible that Mp10 and EDS1 act in a mutually repressive manner or could inversely regulate distinct OG-inducible DTI pathways, activating one while suppressing another. Given the likelihood of multiple, redundant OG perception and signalling complexes (Gravino *et al*., 2017; Blaschek, 2024), EDS1 absence may be compensated by alternative OG receptors operating independently of EDS1 in the presence of Mp10, enhancing plant resilience. Mp10 activity may be influenced by its interactions with AMSH proteins (Gravino *et al*., 2024), whose functions could vary depending on the presence or absence of EDS1. Nevertheless, our findings identify EDS1 as a key player in Mp10 dual role: suppressing DTI/PTI responses while also eliciting ETI.

Aphids are highly specialised feeders, and their feeding actions affect only a few cells in close proximity to their stylets (Tjallingii & Esch, 1993). This localised feeding strategy is evidenced by minimal calcium release upon aphid stylet penetrations (Vincent *et al*., 2017; Joyce, 2023) compared to the robust calcium release triggered by piercing-sucking thrips (Joyce, 2023) or leaf-chewing caterpillars (Toyota *et al*., 2018). Moreover, aphids deliver small quantities of effectors into plant cells, possibly in a sequential manner and tailored to specific cell types (Sanchez-Garrido *et al*., 2022). In contrast, experimental applications of OGs and Mp10 affect entire leaves or plants. Thus, the impacts of OGs and Mp10 are likely to be more nuanced and temporally refined within the aphid-feeding site, where additional factors shape the final outcomes of plant-aphid interactions. In line with this is the finding that Mp10 is present in the acrostyle at the tip of the stylets from where it is likely to be delivered into the cell cytoplasm during the brief salivation periods in the probing phase (Brault *et al*., 2010; Deshoux *et al*., 2022), as evidenced by the presence of Mp10 in the cytoplasm of cells near aphid stylets (Mugford *et al*., 2016). Therefore, Mp10 is likely to be present at the early feeding stages, at the time when aphids just have started their probing behaviour, when suppression of DTI/PTI responses is needed. Aphids also deliver other effectors, including cathepsin B-like proteins (CathBs) such as CathB6, which interacts with EDS1, recruits it to processing bodies, and suppresses EDS1-mediated immunity (Liu *et al*., 2024).

This study, together with previous research on aphid effectors and plant responses, contributes to a comprehensive model encompassing all key players involved (Fig. 8). These findings underscore the dynamic interplay between plant defences and aphid countermeasures, highlighting the critical role of OGs in activating DTI and the adaptive strategies aphids employ, such as suppressing OG production and deploying effectors like Mp10, to modulate host defences. This knowledge will contribute to devising durable and sustainable strategies to control aphids and potentially other sap-sucking insects and the viruses they transmit.

## ACKNOWLEDGMENTS

We thank Jen Sheen (Massachusetts General Hospital and Harvard Medical School) for providing seeds of Arabidopsis *cpk5 cpk6 cpk11*; Cyril Zipfel (University of Zurich and The Sainsbury Laboratory) for providing seeds of Arabidopsis *bak1-5 bkk1-1*; Jonathan Jones (The Sainsbury Laboratory) for providing seeds of Arabidopsis *eds1-2*, and *N. benthamiana eds1* and NahG; Sophien Kamoun (The Sainsbury Laboratory) for providing the pCB301-p19 plasmid and seeds of *N. benthamiana nrg1-1* and *nrg1-2*; Phil Carella (John Innes Centre) for providing seeds of *N. benthamiana adr1 nrg1*; Silke Robatzek (LMU Munich Biocenter) for providing the AtFLS2-eGFP construct; Phil Robinson (John Innes Centre) for photography of Arabidopsis and *N. benthamiana*. We are grateful to the John Innes Centre (JIC) Horticultural Services for growing plants and the JIC Entomology

Facility for rearing aphid stocks. This work was supported by the Biotechnology and Biological Sciences Research Council (BBSRC) grants (grant numbers: BB/R009481/1, BB/V008544/1, BBS/E/J/000PR9795, BBS/E/J/000PR9796, and BBS/E/J/000PR9797), and the John Innes Foundation funded to SAH. Additional support was received from the Ministero dell’Università e della Ricerca (MUR) PRIN 2022 and the Sapienza University of Rome Medi_Progetti_2023 (grant numbers: PRIN2022WLZ4HB and RM123188F70CF274) funded to GDL, and the EMBO Short-Term Fellowship (ASTF number: 477-2016) and the Sapienza University of Rome Progetti di avvio alla ricerca 2016 (project number: AR21615506343970) funded to MG.

## AUTHOR CONTRIBUTIONS

MG, GDL, and SAH acquired funding for the research. SAH managed the project and staff. MG, STM, DP, JJ, FC, GDL, and SAH planned and designed research. MG, STM, DP, and JJ performed experiments. MG, STM, DP, and JJ collected and analysed data. MG, STM, DP, JJ, FC, GDL, and SAH interpreted data. MG, STM, CD, and DCP contributed new plant material. MG, DP, FC, GDL, and SAH wrote the original draft of the paper. All authors reviewed, edited, and approved the final manuscript.

## DATA AVAILABILITY

The data that support the findings of this study are available from the corresponding author upon reasonable request.

The following Supporting Information is available for this article:

**Fig. S1** Effect of β-oestradiol on the activation of the OG machine (OGM) and on aphid fecundity

**Fig. S2** Genotyping of the *brassinosteroid-insensitive 1 (BRI1)-associated receptor kinase 1* (*BAK1*) allele *bak1-5*

**Fig. S3** Content of oligogalacturonides (OGs) in aphid-infested and control leaves

**Fig. S4** High-performance anion-exchange chromatography (HPAEC) with pulsed amperometric detector (PAD) profiles of oligogalacturonides (OGs) extracted with or without sodium sulphite from aphid-infested and control leaves

**Fig. S5** Characterisation of *Mp10* transcript levels in response to dexamethasone (DEX) in Arabidopsis DEX-inducible *Mp10* transgenic lines

**Fig. S6** The aphid effector Mp10 specifically suppresses reactive oxygen species (ROS) bursts triggered by bacterial flg22, but not fungal chitin

**Fig. S7** The aphid effector Mp10 promotes EDS1-dependent defense activation in *Arabidopsis thaliana*

**Table S1** List of primer sequences used in this study

**Table S2** *Arabidopsis thaliana* genes and elements involved in both oligogalacturonide (OG) signalling and immunity to *Myzus persicae*

**Table S3** Post hoc comparisons - genotype ✻ treatment, relative to Fig. 6b

**Table S4** List of icons, including attribution and license, used to generate the model in Fig. 8

---

**Fig. S1.**
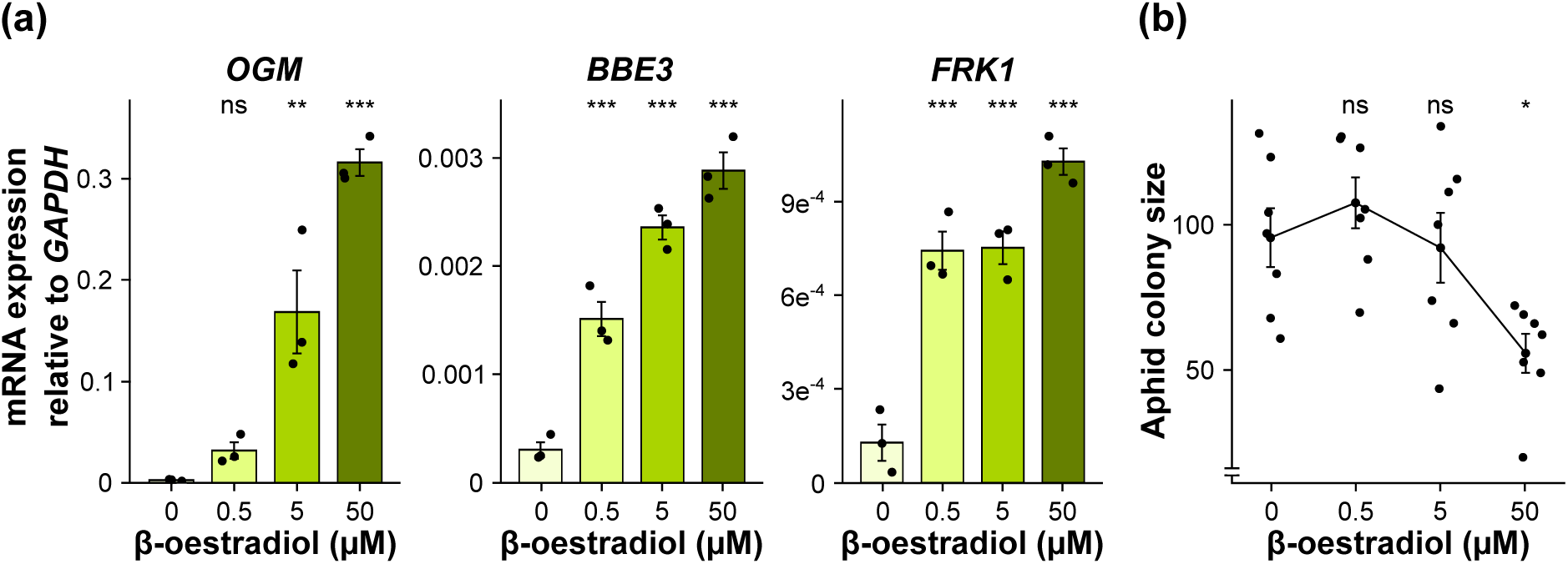
Effect of β-oestradiol on the activation of the OG machine (OGM) and on aphid fecundity. (a) One leaf from 4-week (w)-old Arabidopsis *XVE:OGM* transgenic plants was infiltrated with dimethyl sulfoxide (DMSO) or 0.5, 5, or 50 μg ml^-1^ β-oestradiol (dissolved in DMSO) (x-axis). Transcript levels of *OGM*, *BERBERINE BRIDGE ENZYME-LIKE 3* (*BBE3*/*RetOx*/*FOX1*), and *FLG22-INDUCED RECEPTOR-LIKE KINASE 1* (*FRK1*) relative to *GLYCERALDEHYDE 3-PHOSPHATE DEHYDROGENASE* (*GAPDH*) (y-axis) were measured by quantitative reverse transcriptase (qRT)-PCR 3 days post infiltration (dpi). Bars are means ± SEM. n = 3 replicates. (b) One leaf from 4-w-old Arabidopsis Col-0 WT was infiltrated with DMSO or 0.5, 5, or 50 μg ml^-1^ β-oestradiol (dissolved in DMSO) (x-axis). 3 dpi, one 6-d-old asexually reproducing *M. persicae* adult female was caged on the infiltrated leaf. After 10 d, the number of individual aphids (adults + nymphs) within the cage was counted to obtain the aphid colony size (y-axis). Results are shown as means ± SEM. n = 8 replicates. (a, b) Asterisks indicate significant differences between DMSO- and β-oestradiol- treated samples as determined by one-way ANOVA with post-hoc Tukey HSD test (*, *P* < 0.05; **, *P* < 0.01; ***, *P* < 0.001; ns, not significant).

**Fig. S2.**
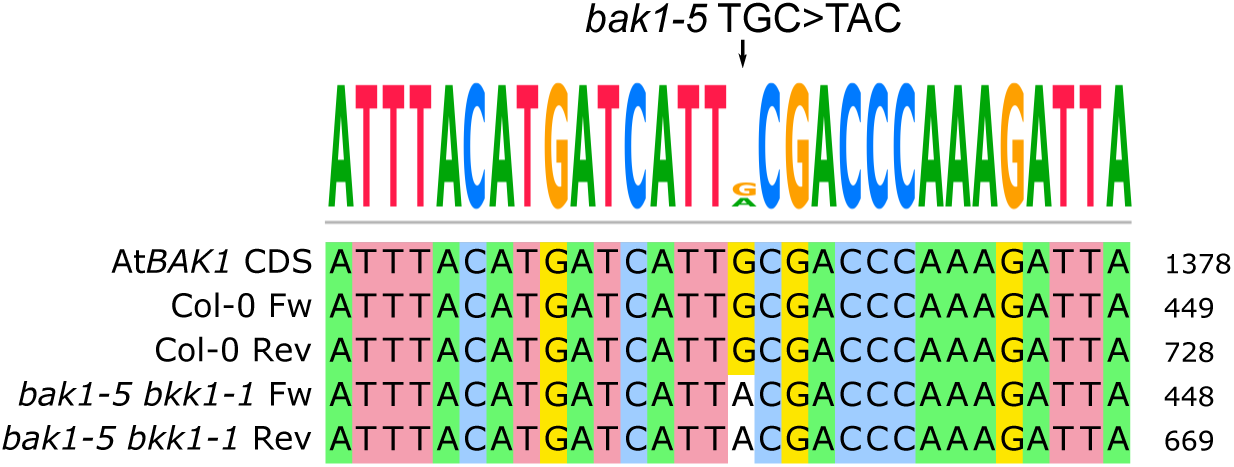
Genotyping of the *brassinosteroid-insensitive 1 (BRI1)-associated receptor kinase 1* (*BAK1*) allele *bak1-5*. Point mutation TGC>TAC (indicated by the arrow on top of the graph) in the coding DNA sequence (CDS) of transcripts of the *BAK1* allele *bak1-5* was confirmed by sequencing the reverse transcriptase (RT)-PCR products of Col-0 wild type (WT) and *bak1-5 bkk1-1* mutant plants, obtained using gene-specific primers listed in Supporting Information Table S2 and complementary DNA (cDNA) as template.

**Fig. S3.**
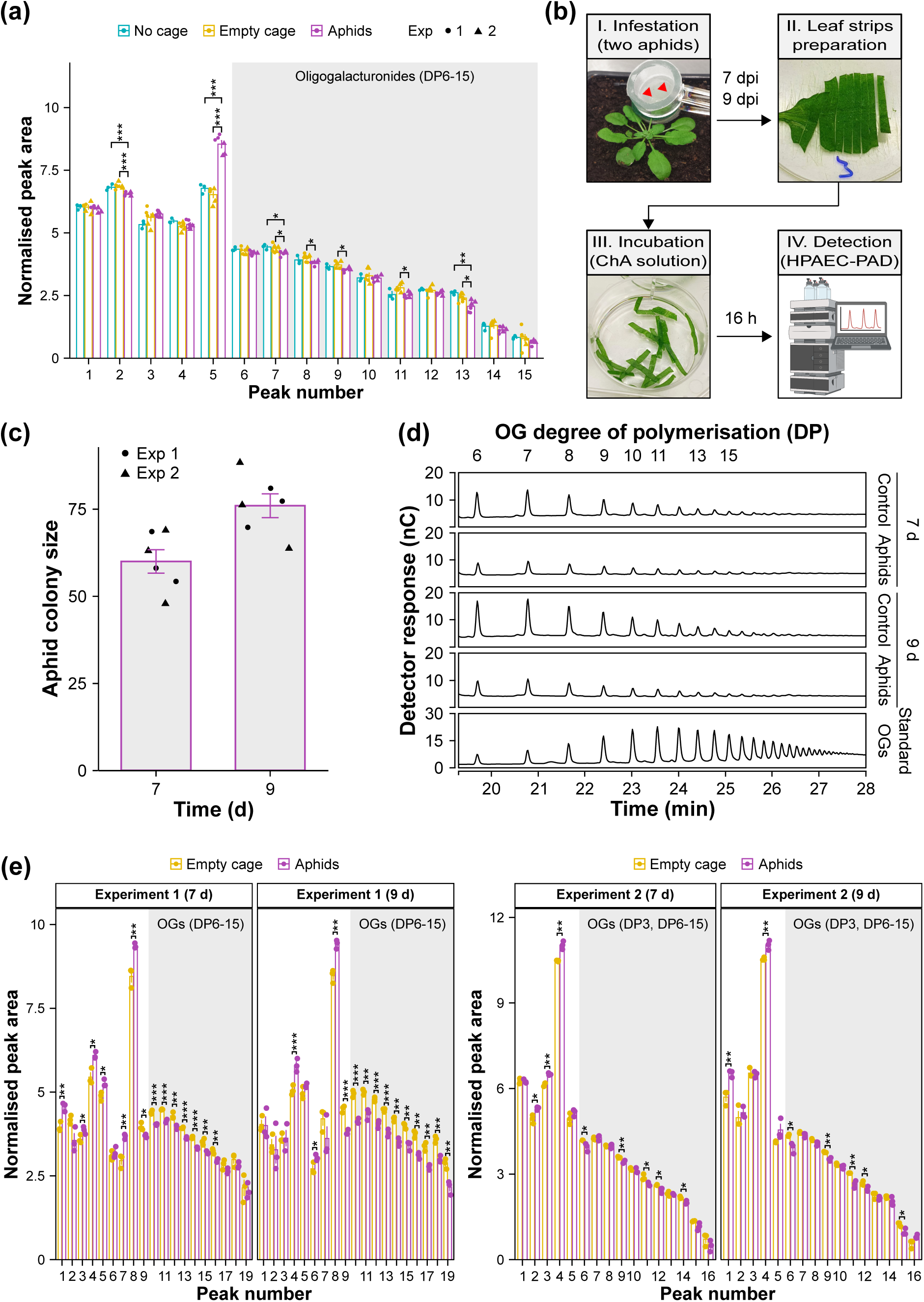
Content oligogalacturonides (OGs) in aphid-infested and control leaves. (a) Content of common peaks, including different OG oligomers (DP6-15), in the chromatograms of controls and aphid-infested leaves for 6 h. The x-axis shows the peak number: peaks 1 to 5 (not highlighted) cannot be clearly identified as pectic fragments; peaks 6 to 15 (highlighted in grey) correspond to OG DP6 to 15. The y-axis shows the peak area (nC min), normalised using the compositional data normalisation (CoDA) method. Bars are means ± SEM. Observations are indicated with dots for experiment (exp) 1 or triangles for exp 2. n = 3 replicates in each experiment. (b) Experiment design to detect OGs during late aphid feeding. Leaves from 4-week-old Arabidopsis Col-0 wild-type (WT) plants were caged with two 6-d-old asexually reproducing *M. persicae* adult females (panel I) or without aphids (empty cage), as a control. After 7 or 9 days post infestation (dpi), leaves were excised and, after removing all aphids, sliced into strips (panel II). Leaf strips were incubated with strong chelating agents supplemented with sodium sulphite (ChA solution) for 16 h (panel III). The leaf diffusates in incubation medium were analysed by high-performance anion-exchange chromatography (HPAEC) with a pulsed amperometric detector (PAD) (panel IV). The illustration in panel IV was created with BioRender.com (agreement number: EI25PBPRUZ). (c) Aphid colony size (y-axis) inside clip cages after 7 and 9 dpi (x-axis). Bars are means ± SEM. Observations are indicated with dots for experiment (exp) 1 or triangles for exp 2. n = 3 replicates in each experiment. (d) Chromatographic analysis of leaf diffusates in ChA solution obtained from leaf strips of plants infested with aphids or non-infested, as a control, for 7 and 9 d. The numbers on top of the graph indicate the degree of polymerisation (DP) of different OG oligomers and refer to the corresponding peaks below. The x-axis shows the retention time measured in min. The y-axis shows the detector response measured in nanocoulombs (nC). The bottom panel shows the profile of 2.5 μg of standard OGs enriched in the DP range 10-15. For each treatment, the result of one representative replicate out of three performed is shown. (e) Content of common peaks, including different OG oligomers (DP3, DP6-15), in the chromatograms of controls and aphid-infested leaves for 7 and 9 d. The x-axis shows the peak number: peaks 1 to 9 and 1 to 5 (not highlighted) cannot be clearly identified as pectic fragments in experiments 1 and 2, respectively; peaks 10 to 19 and 7 to 16 correspond to OG DP6 to 15 in experiments 1 and 2, respectively; peak 6 in experiment 2 corresponds to OG DP3. The y-axis shows the peak area (nC min) of the common peaks, normalised using the compositional data normalisation (CoDA) method. Bars are means ± SEM. Observations are indicated with dots. n = 3 replicates in each experiment. In (a, e), asterisks indicate significant differences between aphid-infested leaves and control samples as determined by one-way ANOVA with post-hoc Tukey HSD test or Student’s *t* test, respectively (*, *P* < 0.05; **, *P* < 0.01; ***, *P* < 0.001).

**Fig. S4.**
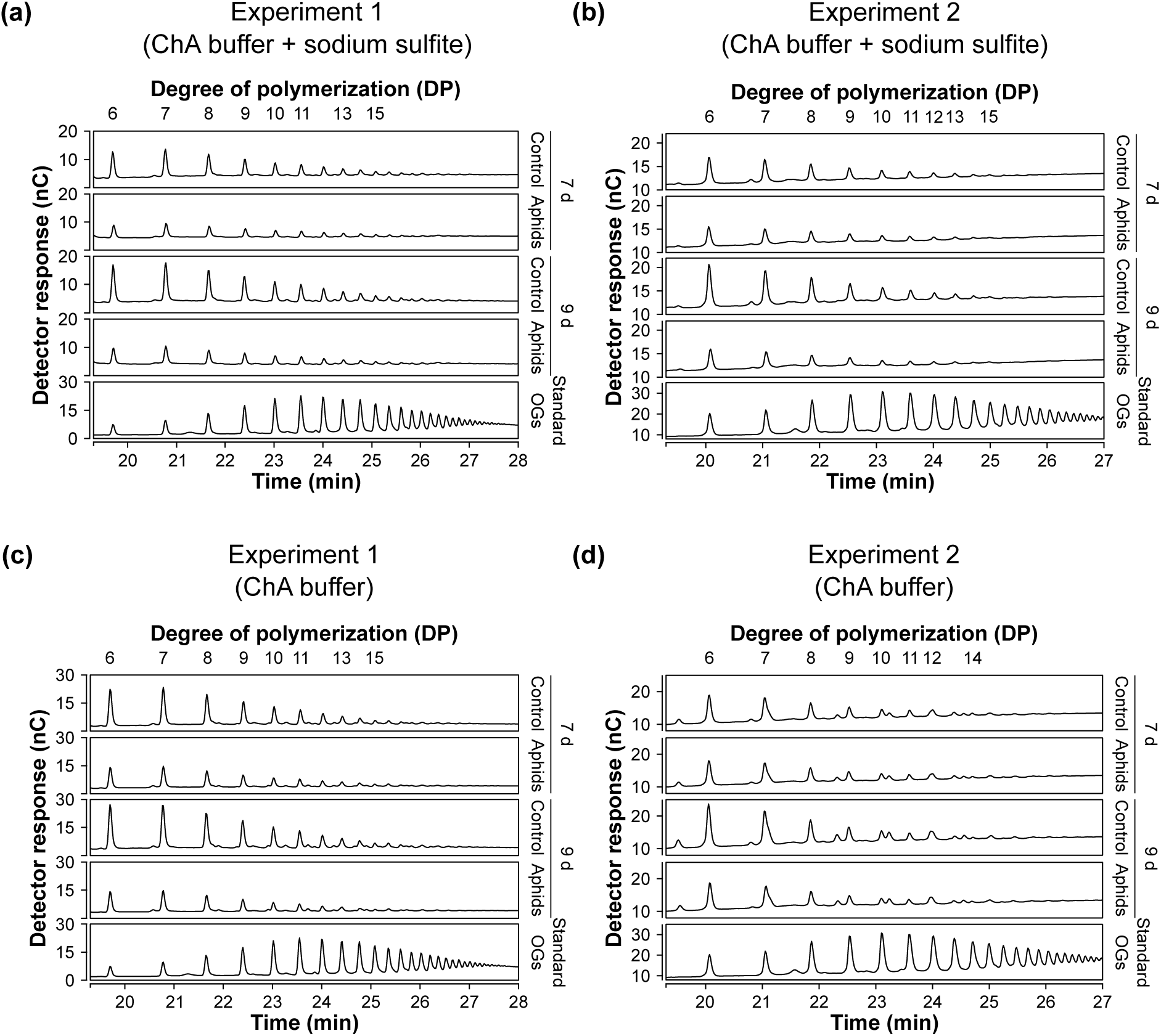
High-performance anion-exchange chromatography (HPAEC) with pulsed amperometric detector (PAD) profiles of oligogalacturonides (OGs) extracted with or without sodium sulphite from aphid-infested and control leaves. Leaf strips were obtained from 4-week-old Arabidopsis Col-0 wild-type (WT) plants infested with aphids or non-infested, as a control, for 7 or 9 days. Leaf strips were incubated for 16 h in strong chelating agents (ChA solution, i.e., 50 mM ammonium acetate pH 5.0, 50 mM CDTA, 50 mM ammonium oxalate) supplemented with 10 mM sodium sulphite (ChA buffer + sodium sulphite), to inhibit the activity of OG oxidases (Benedetti *et al*., 2017) or without sodium sulphite (ChA buffer) to detect eventual presence of oxidised OGs (Benedetti *et al*., 2018). Leaf diffusates in the buffers were then analysed by HPAEC-PAD. (a,b) Experiment 1 and 2, performed with ChA buffer + sodium sulfite. (c,d) Experiment 1 and 2, performed with ChA buffer. (a, b, c, d). The numbers on top of the graphs indicate the degree of polymerisation (DP) of different OG oligomers and refer to the corresponding peaks below. The x-axis shows the retention time for different OG oligomers measured in min. The y-axis shows the detector response for different OG oligomers measured in nanocoulombs (nC). The bottom panels show the profile of 2.5 μg of standard OGs enriched in the DP range 10-15. For each experiment, the result of one representative replicate out of three performed is shown.

**Fig. S5.**
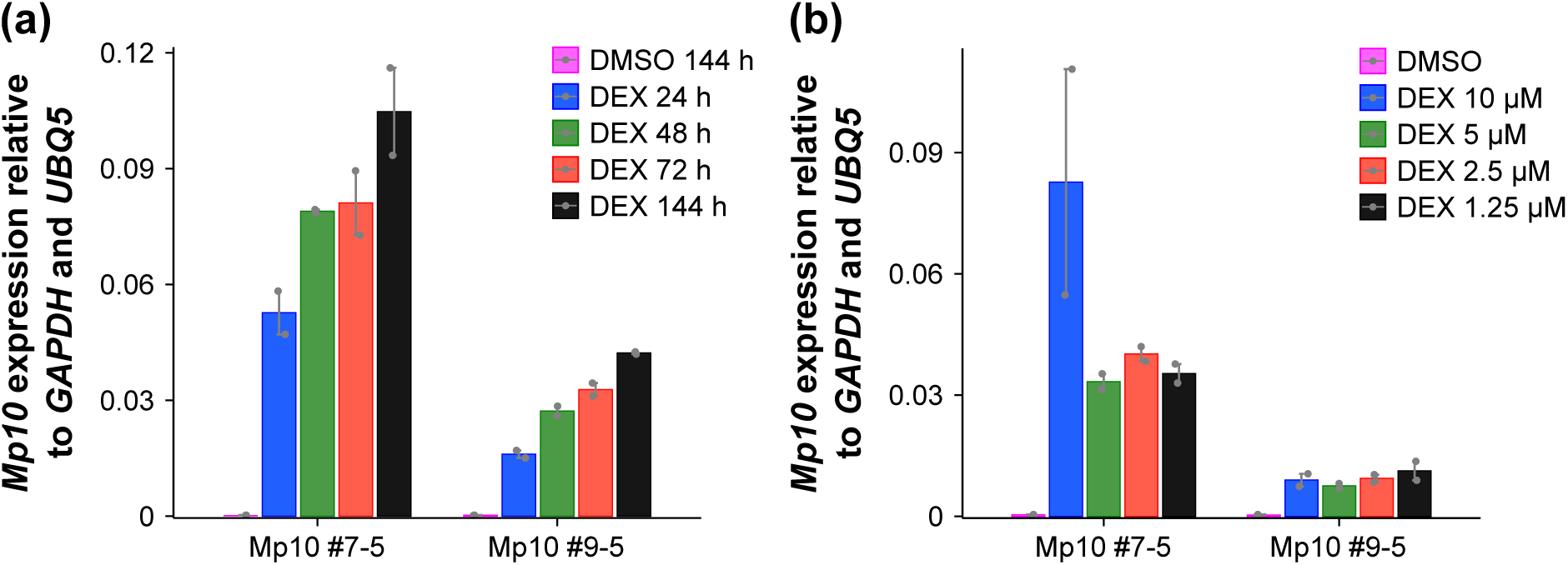
Characterisation of *Mp10* transcript levels in response to dexamethasone (DEX) in Arabidopsis DEX-inducible *Mp10* transgenic lines. Transcript levels of *Mp10* relative to *GLYCERALDEHYDE 3-PHOSPHATE DEHYDROGENASE* (*GAPDH*) and *UBIQUITIN 5* (*UBQ5*) (y-axis) were measured by quantitative reverse transcriptase (qRT)-PCR in leaves of 4-week-old Arabidopsis *Mp10* transgenic lines (#7-5 and #9-5) (x-axis) at 24, 48, 72, and 144 h after spraying with 20 μM DEX and at 144 h after spraying with dimethyl sulfoxide (DMSO), as a control (a), and at 72 h after spraying with DMSO or 10, 5, 2.5, or 1.25 μM DEX (b). Results are shown as mean ± SEM. n = 2 biological repeats.

**Fig. S6.**
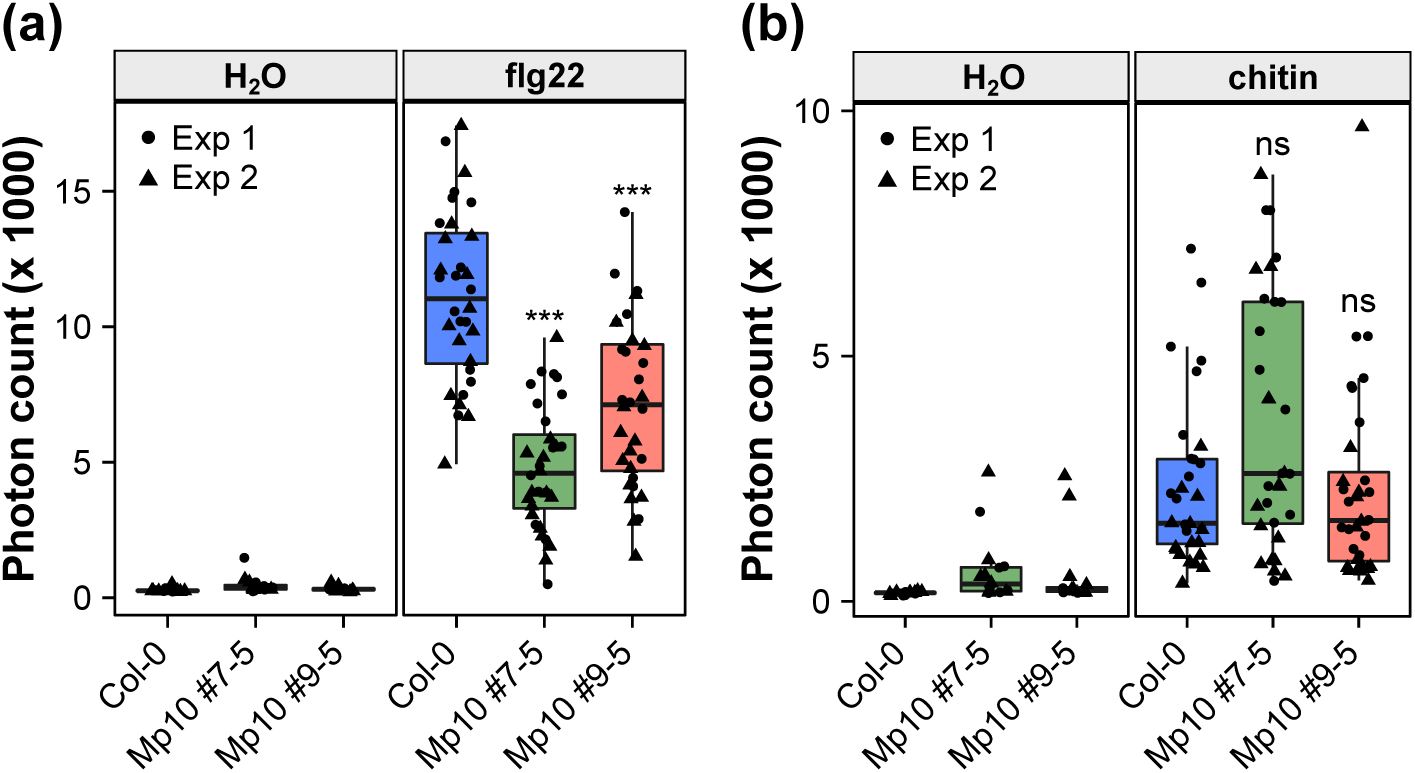
The aphid effector Mp10 specifically suppresses reactive oxygen species (ROS) bursts triggered by bacterial flg22, but not fungal chitin. (a) Total ROS production over a period of 8 h after flg22 treatment. (b) Total ROS production over a period of 5 h after chitin treatment. In (a, b), 3- to 4-week (w)-old plants of *Arabidopsis thaliana* Col-0 wild type (WT) and dexamethasone (DEX)-inducible Mp10 transgenic lines (#7-5 and #9-5) were sprayed with 1.25 μM DEX and, 3 d later, treated with H_2_O, 100 nM flg22, or 500 μg ml^-1^ chitin, dissolved in water. The y-axis shows the total photon count. The x-axis shows the plant genotype. Boxplots show the median, the 25^th^ and 75^th^ percentiles, the most extreme data points (whiskers’ extensions), and the observations as black-filled circles for experiment (exp) 1 or triangles for exp 2. *n* = 8 and 16 leaf discs treated with H_2_O and flg22/chitin, respectively, in each experiment. Asterisks indicate significant differences between Col-0 WT and Mp10 transgenic lines treated with flg22 or chitin, as determined by one-way ANOVA with post-hoc Tukey HSD test (***, *P* < 0.001; ns, not significant).

**Fig. S7.**
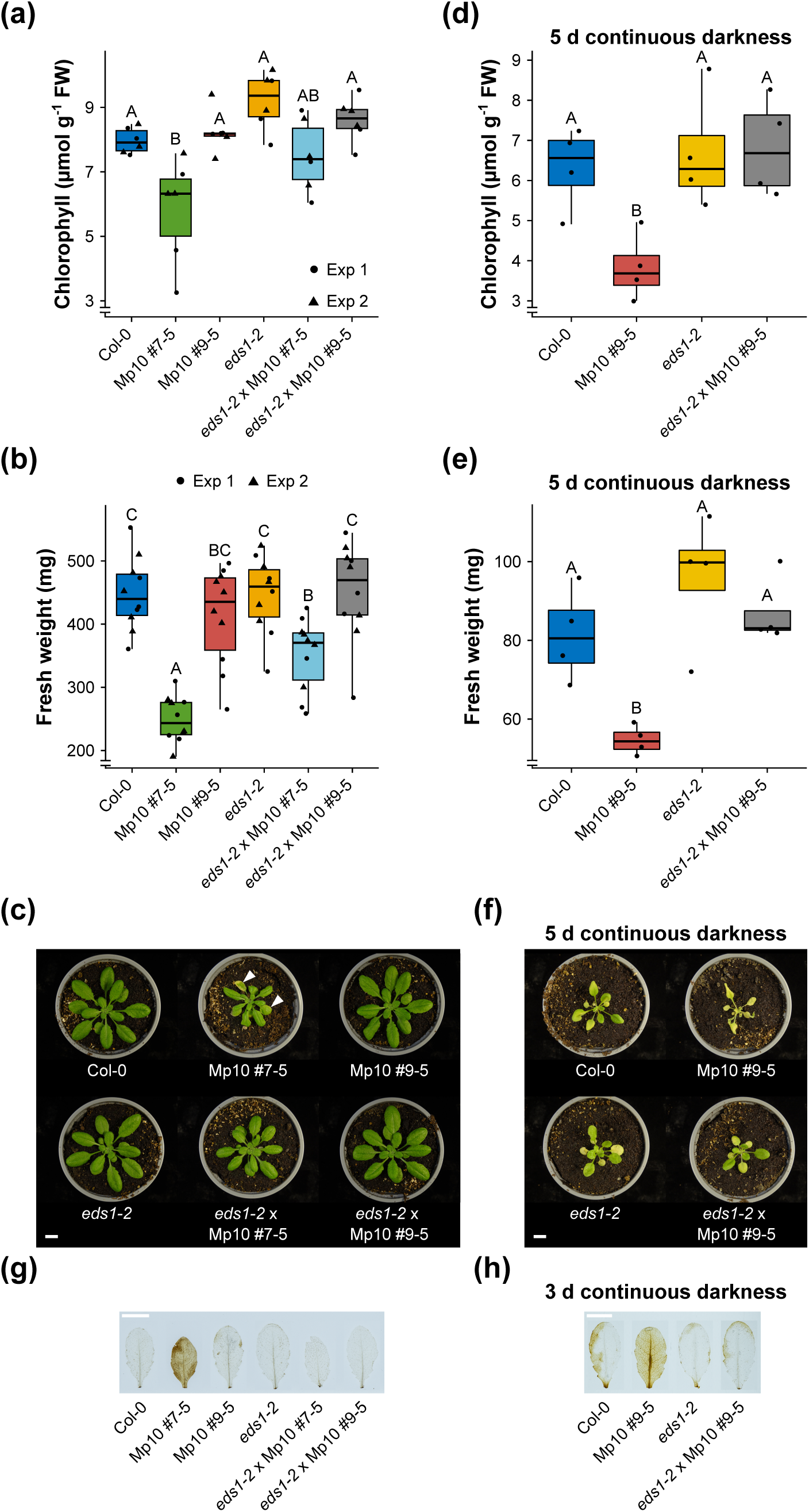
The aphid effector Mp10 promotes EDS1-dependent defense activation in *Arabidopsis thaliana*. (a, b, c, d, e, f) Chlorophyll and fresh weight measurements in Arabidopsis plants pretreated with 1.25 μM dexamethasone (DEX) and grown under normal conditions (a, b, c) or for 5 d under continuous darkness (d, e, f). In (a, d), the y-axis shows the micromolar (μmol) chlorophyll content per gram (g) of fresh weight (FW). In (b, e), the y- axis shows the fresh weight measured in mg. (c) Representative pictures showing plant size as well as symptoms of chlorosis and senescence, as indicated by arrowheads, only in the Arabidopsis Mp10 #7-5 line but not in other plants. (f) Representative pictures showing plant size as well as increased symptoms of chlorosis and senescence in Arabidopsis Mp10 #9- 5 line compared to Col-0, *eds1-2*, and *eds1-2* x Mp10 #9-5. In (a, b, d, e), the x-axis shows the plant genotype. Boxplots show the median, the 25^th^ and 75^th^ percentiles, the most extreme data points (whiskers’ extensions), and the observations as black-filled circles for experiment (exp) 1 or triangles for exp 2. n ≥ 3 samples in each experiment. Different letters indicate significant differences between samples as determined by one-way ANOVA with post-hoc Tukey HSD test (*P* < 0.001, *P* < 0.001, *P* < 0.05, *P* < 0.01, respectively). (g, h) *In situ* detection of hydrogen peroxide by DAB staining in the indicated plant genotypes pretreated with DEX and grown under normal conditions (g) or for 3 d under continuous darkness (h). In (c, f, g, h), scale bars = 1 cm.

**Table S1.**
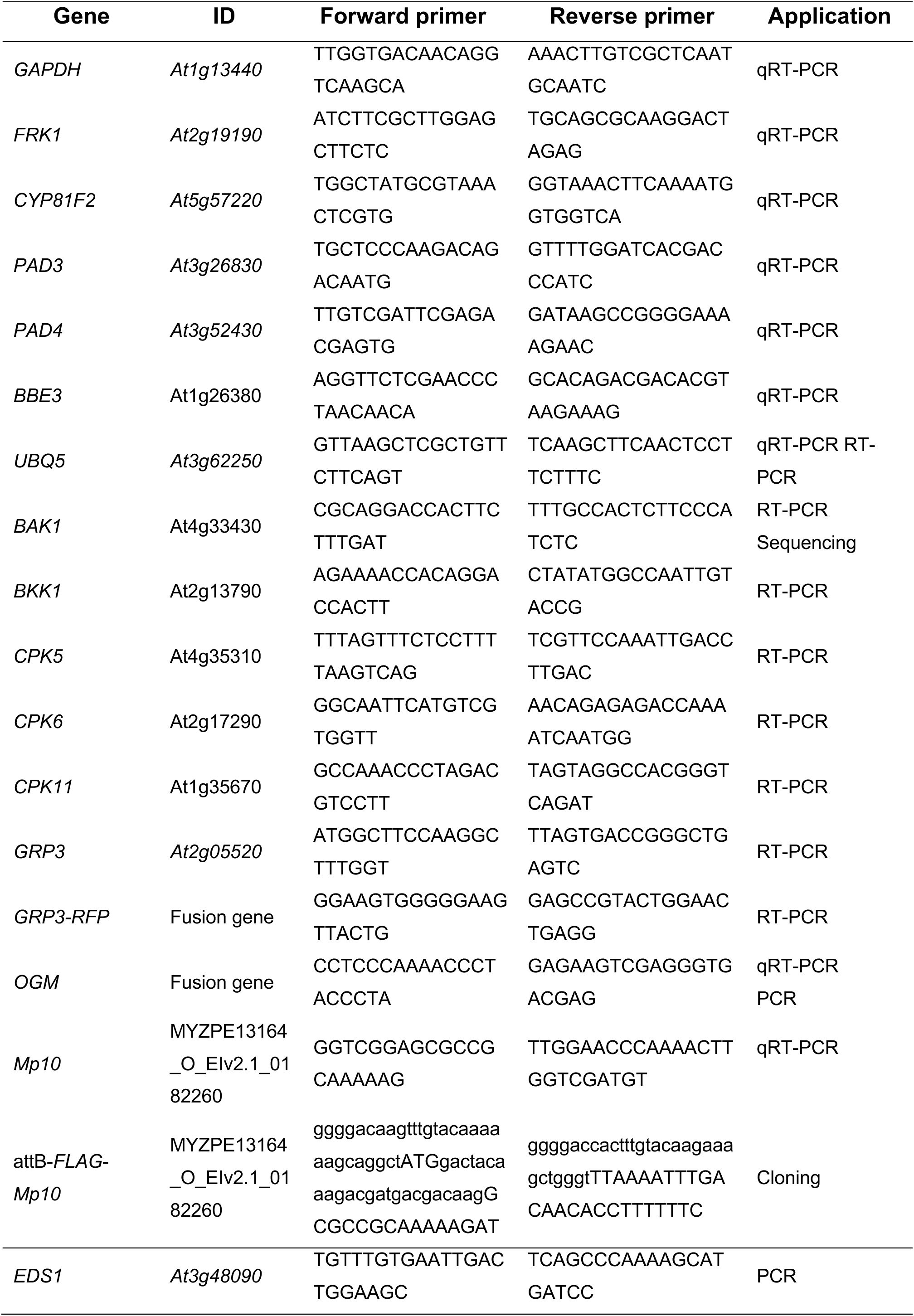
List of primer sequences used in this study.

**Table S2.**
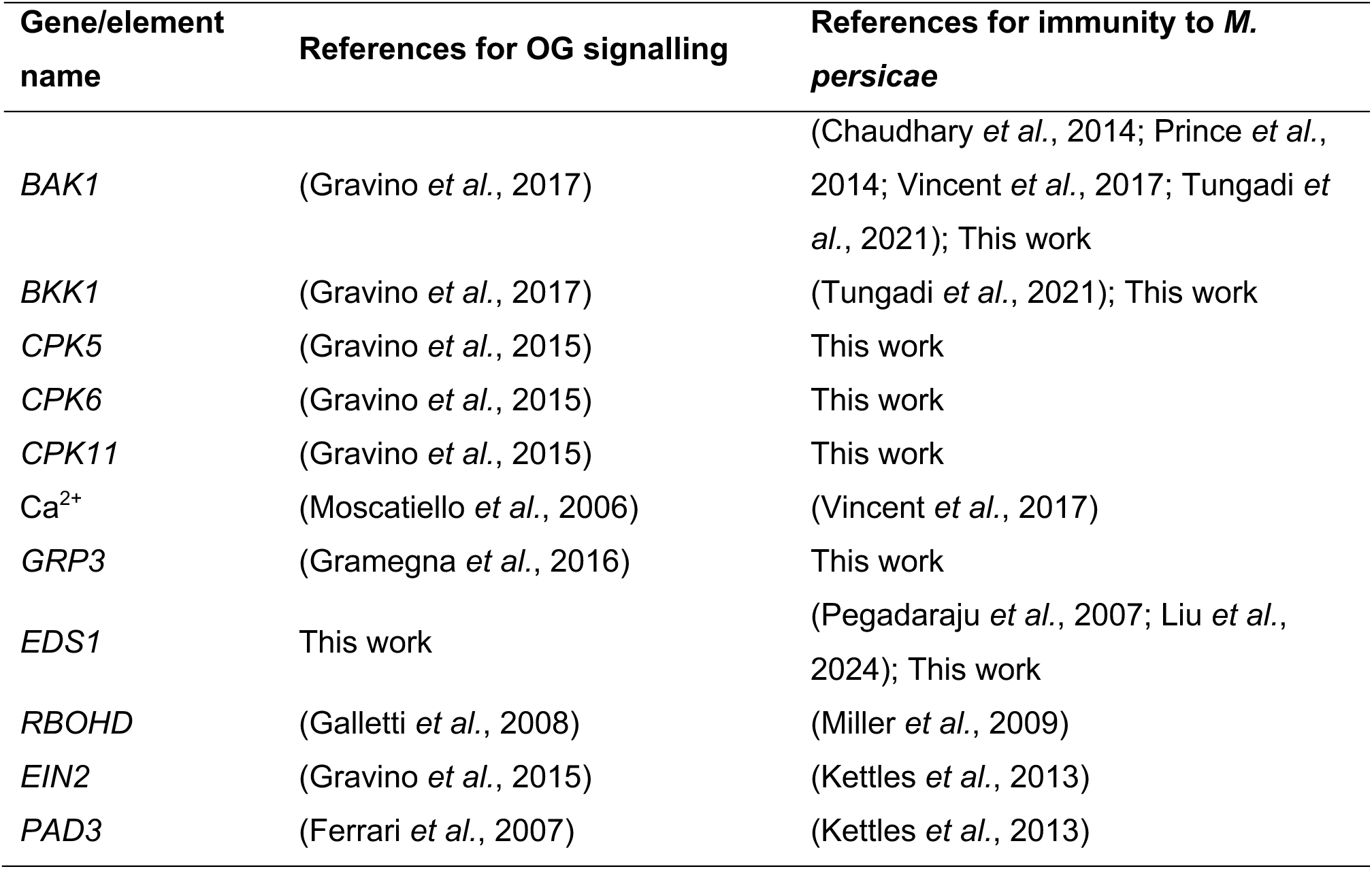
*Arabidopsis thaliana* genes and elements involved in both oligogalacturonide (OG) signalling and immunity to *Myzus persicae*.

**Table S3.** Post hoc comparisons - genotype ✻ treatment, relative to Fig. 6b.

|  |  | **Mean**<br>**Difference** | **SE** | **df** | **t** | **p<sub>tukey</sub>** | **Significance** |
| --- | --- | --- | --- | --- | --- | --- | --- |
| (Col-0 H<sub>2</sub>O) | (Mp10 #7-5 H<sub>2</sub>O) | 25.187 | 6.342 | 180 | 3.971 | 0.006 | ** |
|  | (Mp10 #9-5 H<sub>2</sub>O) | 25.875 | 6.342 | 180 | 4.08 | 0.004 | ** |
|  | (eds1-2 H<sub>2</sub>O) | 0.937 | 6.342 | 180 | 0.148 | 1 |  |
|  | (eds1-2 x Mp10 #7-5 H<sub>2</sub>O) | -17.625 | 6.342 | 180 | -2.779 | 0.198 |  |
|  | (eds1-2 x Mp10 #9-5 H<sub>2</sub>O) | -15.75 | 6.342 | 180 | -2.483 | 0.357 |  |
|  | (Col-0 OG) | 30.562 | 6.342 | 180 | 4.819 | < .001 | *** |
|  | (Mp10 #7-5 OG) | 25.438 | 6.342 | 180 | 4.011 | 0.005 | ** |
|  | (Mp10 #9-5 OG) | 27.125 | 6.342 | 180 | 4.277 | 0.002 | ** |
|  | (eds1-2 OG) | 0.437 | 6.342 | 180 | 0.069 | 1 |  |
|  | (eds1-2 x Mp10 #7-5 OG) | 12.25 | 6.342 | 180 | 1.931 | 0.738 |  |
|  | (eds1-2 x Mp10 #9-5 OG) | 11.25 | 6.342 | 180 | 1.774 | 0.83 |  |
| (Mp10 #7-5 H<sub>2</sub>O) | (Mp10 #9-5 H<sub>2</sub>O) | 0.687 | 6.342 | 180 | 0.108 | 1 |  |
|  | (eds1-2 H<sub>2</sub>O) | -24.25 | 6.342 | 180 | -3.824 | 0.01 | ** |
|  | (eds1-2 x Mp10 #7-5 H<sub>2</sub>O) | -42.813 | 6.342 | 180 | -6.75 | < .001 | *** |
|  | (eds1-2 x Mp10 #9-5 H<sub>2</sub>O) | -40.938 | 6.342 | 180 | -6.455 | < .001 | *** |
|  | (Col-0 OG) | 5.375 | 6.342 | 180 | 0.847 | 0.999 |  |
|  | (Mp10 #7-5 OG) | 0.25 | 6.342 | 180 | 0.039 | 1 |  |
|  | (Mp10 #9-5 OG) | 1.937 | 6.342 | 180 | 0.305 | 1 |  |
|  | (eds1-2 OG) | -24.75 | 6.342 | 180 | -3.902 | 0.007 | ** |
|  | (eds1-2 x Mp10 #7-5 OG) | -12.938 | 6.342 | 180 | -2.04 | 0.666 |  |
|  | (eds1-2 x Mp10 #9-5 OG) | -13.938 | 6.342 | 180 | -2.198 | 0.554 |  |
| (Mp10 #9-5 H<sub>2</sub>O) | (eds1-2 H<sub>2</sub>O) | -24.938 | 6.342 | 180 | -3.932 | 0.007 | ** |
|  | (eds1-2 x Mp10 #7-5 H<sub>2</sub>O) | -43.5 | 6.342 | 180 | -6.859 | < .001 | *** |
|  | (eds1-2 x Mp10 #9-5 H<sub>2</sub>O) | -41.625 | 6.342 | 180 | -6.563 | < .001 | *** |
|  | (Col-0 OG) | 4.688 | 6.342 | 180 | 0.739 | 1 |  |
|  | (Mp10 #7-5 OG) | -0.437 | 6.342 | 180 | -0.069 | 1 |  |
|  | (Mp10 #9-5 OG) | 1.25 | 6.342 | 180 | 0.197 | 1 |  |
|  | (eds1-2 OG) | -25.438 | 6.342 | 180 | -4.011 | 0.005 | ** |
|  | (eds1-2 x Mp10 #7-5 OG) | -13.625 | 6.342 | 180 | -2.148 | 0.589 |  |
|  | (eds1-2 x Mp10 #9-5 OG) | -14.625 | 6.342 | 180 | -2.306 | 0.476 |  |
| (eds1-2 H<sub>2</sub>O) | (eds1-2 x Mp10 #7-5 H<sub>2</sub>O) | -18.562 | 6.342 | 180 | -2.927 | 0.141 |  |
|  | (eds1-2 x Mp10 #9-5 H<sub>2</sub>O) | -16.687 | 6.342 | 180 | -2.631 | 0.271 |  |
|  | (Col-0 OG) | 29.625 | 6.342 | 180 | 4.671 | < .001 | *** |
|  | (Mp10 #7-5 OG) | 24.5 | 6.342 | 180 | 3.863 | 0.008 | ** |
|  | (Mp10 #9-5 OG) | 26.188 | 6.342 | 180 | 4.129 | 0.003 | ** |
|  | (eds1-2 OG) | -0.5 | 6.342 | 180 | -0.079 | 1 |  |
|  | (eds1-2 x Mp10 #7-5 OG) | 11.313 | 6.342 | 180 | 1.784 | 0.825 |  |
|  | (eds1-2 x Mp10 #9-5 OG) | 10.313 | 6.342 | 180 | 1.626 | 0.897 |  |
| (eds1-2 x Mp10 #7-5 H<sub>2</sub>O) | (eds1-2 x Mp10 #9-5 H<sub>2</sub>O) | 1.875 | 6.342 | 180 | 0.296 | 1 |  |
|  | (Col-0 OG) | 48.188 | 6.342 | 180 | 7.598 | < .001 | *** |
|  | (Mp10 #7-5 OG) | 43.063 | 6.342 | 180 | 6.79 | < .001 | *** |
|  | (Mp10 #9-5 OG) | 44.75 | 6.342 | 180 | 7.056 | < .001 | *** |
|  | (eds1-2 OG) | 18.062 | 6.342 | 180 | 2.848 | 0.169 |  |
|  | (eds1-2 x Mp10 #7-5 OG) | 29.875 | 6.342 | 180 | 4.71 | < .001 | *** |
|  | (eds1-2 x Mp10 #9-5 OG) | 28.875 | 6.342 | 180 | 4.553 | < .001 | *** |
| (eds1-2 x Mp10 #9-5 H<sub>2</sub>O) | (Col-0 OG) | 46.313 | 6.342 | 180 | 7.302 | < .001 | *** |
|  | (Mp10 #7-5 OG) | 41.188 | 6.342 | 180 | 6.494 | < .001 | *** |
|  | (Mp10 #9-5 OG) | 42.875 | 6.342 | 180 | 6.76 | < .001 | *** |
|  | (eds1-2 OG) | 16.187 | 6.342 | 180 | 2.552 | 0.315 |  |
|  | (eds1-2 x Mp10 #7-5 OG) | 28 | 6.342 | 180 | 4.415 | 0.001 | ** |
|  | (eds1-2 x Mp10 #9-5 OG) | 27 | 6.342 | 180 | 4.257 | 0.002 | ** |
| (Col-0 OG) | (Mp10 #7-5 OG) | -5.125 | 6.342 | 180 | -0.808 | 1 |  |
|  | (Mp10 #9-5 OG) | -3.438 | 6.342 | 180 | -0.542 | 1 |  |
|  | (eds1-2 OG) | -30.125 | 6.342 | 180 | -4.75 | < .001 | *** |
|  | (eds1-2 x Mp10 #7-5 OG) | -18.313 | 6.342 | 180 | -2.887 | 0.154 |  |
|  | (eds1-2 x Mp10 #9-5 OG) | -19.313 | 6.342 | 180 | -3.045 | 0.104 |  |
| (Mp10 #7-5 OG) | (Mp10 #9-5 OG) | 1.687 | 6.342 | 180 | 0.266 | 1 |  |
|  | (eds1-2 OG) | -25 | 6.342 | 180 | -3.942 | 0.006 | ** |
|  | (eds1-2 x Mp10 #7-5 OG) | -13.188 | 6.342 | 180 | -2.079 | 0.638 |  |
|  | (eds1-2 x Mp10 #9-5 OG) | -14.188 | 6.342 | 180 | -2.237 | 0.525 |  |
| (Mp10 #9-5 OG) | (eds1-2 OG) | -26.688 | 6.342 | 180 | -4.208 | 0.002 | ** |
|  | (eds1-2 x Mp10 #7-5 OG) | -14.875 | 6.342 | 180 | -2.345 | 0.449 |  |
|  | (eds1-2 x Mp10 #9-5 OG) | -15.875 | 6.342 | 180 | -2.503 | 0.345 |  |
| (eds1-2 OG) | (eds1-2 x Mp10 #7-5 OG) | 11.813 | 6.342 | 180 | 1.862 | 0.78 |  |
|  | (eds1-2 x Mp10 #9-5 OG) | 10.813 | 6.342 | 180 | 1.705 | 0.864 |  |
| (eds1-2 x Mp10 #7-5 OG) | (eds1-2 x Mp10 #9-5 OG) | -1 | 6.342 | 180 | -0.158 | 1 |  |

\*\* p < .01, \*\*\* p < .001
*Note.* P-value adjusted for comparing a family of 12 estimates.

**Table S4.** List of icons, including attribution and license, used to generate the model in Fig. 8.

| Icon | Attribution | License |
| --- | --- | --- |
| aphid | DBCLS | CC-BY 4.0 Unported |
|  | https://togotv.dbcls.jp/en/pics.html | https://creativecommons.org/licenses/by/4.0/ |
| cell_group | JhonnyXC | CC0 |
|  | https://github.com/JRider16 | https://creativecommons.org/publicdomain/zero/1.0/ |
| channel-membrane-red-2 | Servier https://smart.servier.com/ | CC-BY 3.0 Unported |
|  |  | https://creativecommons.org/licenses/by/3.0/ |
| Lightning bolt | https://www.vecteezy.com/free-vector/lightning-bolt>Lightning Bolt Vectors by Vecteezy | N/A |

*Note.* All the material in 4.0 license version was modified.

